# CRISPR-Cas9 Engineered Extracellular Vesicles for the Treatment of Dominant Progressive Hearing Loss

**DOI:** 10.1101/2023.09.14.557853

**Authors:** Xiaoshu Pan, Peixin Huang, Samantha S. Ali, Bryan Renslo, Tarun E Hutchinson, Nina Erwin, Zachary Greenberg, Zuo Ding, Yanjun Li, Athanasia Warnecke, Natalia E. Fernandez, Hinrich Staecker, Mei He

## Abstract

Clinical translation of gene therapy has been challenging, due to limitations in current delivery vehicles such as traditional viral vectors. Herein, we report the use of gRNA:Cas9 ribonucleoprotein (RNP) complexes engineered extracellular vesicles (EVs) for *in vivo* gene therapy. By leveraging a novel high-throughput microfluidic droplet-based electroporation system (μDES), we achieved 10-fold enhancement of loading efficiency and more than 1000-fold increase in processing throughput on loading RNP complexes into EVs (RNP-EVs), compared with conventional bulk electroporation. The flow-through droplets serve as enormous bioreactors for offering millisecond pulsed, low-voltage electroporation in a continuous-flow and scalable manner, which minimizes the Joule heating influence and surface alteration to retain natural EV stability and integrity. In the Shaker-1 mouse model of dominant progressive hearing loss, we demonstrated the effective delivery of RNP-EVs into inner ear hair cells, with a clear reduction of *Myo7a*^*sh1*^ mRNA expression compared to RNP-loaded lipid-like nanoparticles (RNP-LNPs), leading to significant hearing recovery measured by auditory brainstem responses (ABR).

**One sentence summary:** The scalable microfluidic electroporation system enables the loading of gRNA:Cas9 ribonucleoprotein (RNP) complexes into extracellular vesicles, which leads to clinical translation potential employed in hearing disease gene therapy.

---

Genome editing is an emerging and powerful therapeutic tool for treating diverse diseases. Sensorineural hearing loss affects more than 450 million people worldwide and up to 50% of cases has a genetic origin^1-3^. To date, gene delivery to the inner ear is an unmet need and has been largely pursued with engineered adeno-associated viral vectors (AAVs) and cationic lipid nanoparticles (LNPs)^4^. Dominant genetic hearing loss disorders often present progressive hearing loss that would need to be addressed with the genome editing. Using AAVs traditionally could introduce the risk of prolonged expression of endonucleases and off target effects. Additionally, AAVs have a limited loading capacity with cargo size below ∼5Kb, therefore, making them unable to carry larger endonucleases such as gRNA:Cas9 in a single vector system^5^. Although larger lentiviruses could increase cargo capacity up to ∼10Kb, the risk of insertional mutagenesis is still a concern^6^. LNPs-mediated CRISPR gene delivery has been demonstrated in post-mitotic cells such as neurons and hair cells, but the tolerability in the inner ear in vivo is not defined yet^7-9^. Alternatively, extracellular vesicles (EV) based gene delivery is emerging as a safe and highly biocompatible modality for addressing challenges^10-13^ in clinical translation of gene therapy. The recent first-in-human trial using umbilical cord mesenchymal stromal cell derived EVs (hUC-MSC-EVs) demonstrated their potential to attenuate inflammation side effects from cochlear implantation^10,14^. However, EV-mediated delivery of gRNA:Cas9 ribonucleoproteins (RNP) complexes for gene editing in vivo enabling hearing loss recovery has not been explored elsewhere.

Efficiently loading gRNA:Cas9 RNP complexes into EVs has been the major roadblock for developing EV based gene therapy. Current methods for EV cargo loading are in low loading efficiency with low stability^15^, and are unscalable for future clinical translation^16,17^. Utilizing engineered cells as the primary EV producer is limited by cargo type and copy numbers that can be passed to EVs. Therefore, we developed a novel **M**icrofluidic **D**roplet-based EV **E**lectroporation **S**ystem (μDES), which can handle various cargos loaded into EVs in large throughput and high efficiency. The saturated cargo concentration in the confined uniform droplet bioreactors maximizes transfection rate with efficient mass transport. Compared to current existing EV transfection approaches^18-22^, the continuous-flow enables much larger scale in high throughput processing. Only a low-voltage DC power (∼10-60 volts) is needed for this streamlined system, which avoids Joule heating and thermal damage to nanosized EVs^23^. In contrast to chemical transfection which introduces potentially ototoxic chemicals to subsequent *in vivo* applications, the instant low voltage electroporation across flow-through droplets is simple and facile for retaining natural EV property employed in downstream *in vivo* administration. We achieved more than a 10-fold enhancement of loading efficiency and 1000-fold increase in processing throughput on loading gRNA:Cas9 RNP complexes into EVs (RNP-EVs), compared with bulk cuvette electroporation and other transfection methods. We also employed Food and Drug Administration approved pharmaceutical excipient trehalose ^24-28^ in the buffer system to preserve EVs in optimal stability^27,29^ during storage and minimize membrane aggregation^24,25^ or leakage^26^ for future clinical translation.

*Myo7a*, which codes for an unconventional myosin in auditory and vestibular hair cells, plays an essential role in the development of sensory hair cells and signal transduction^30-32^. Mutations in this gene lead to 39–55% of the total cases of Usher syndrome (USH1B), as well as non-syndromic hearing loss dominant and recessive hearing loss (DFNA11 and DFNB2)^33-35^. *Myo7a* variants have also been associated with familial presbycusis^36^. Timely removal of mutant *Myo7a* alleles could prevent the progression of hearing loss in the dominant versions of the disorder, which can be evaluated in the *Myo7a Sh1/WT* Shaker-1 mouse^37^. Therefore, we evaluated our μDES produced RNP-EVs for gene editing in Shaker-1 mouse with the gRNA:Cas9 RNP lipid-like nanoparticle (RNP-LNPs) as the control group. Treated animals exhibited hearing restoration in mice with a single dose administration, and significantly reduced *Myo7a*^*sh1*^ mRNA levels. Editing efficiency was demonstrated by qPCR, Sanger sequencing, and Next Generation Sequencing (NGS) analysis. The μDES produced RNP-EVs displayed high biocompatibility and biodistribution within the inner ear tissue, particularly within outer hair cells (OHCs) and inner hair cells (IHCs). In contrast, the RNP-LNPs control group was mainly distributed into spiral ganglion cells. One of the pathogenic hearing loss mechanisms involved in progressive hearing loss is oxidative stress^38-40^. Immunohistochemical evaluation of the oxidative stress biomarkers 3-Nitrotyrosine (3-NT) and 4-Hydroxy-2-nonenal (4-HNE) supported the functional restoration specifically from sensory hair cells after 6 months with μDES RNP EVs gene editing treatment *in vivo*. No noticeable safety concerns or side effects were observed from treated mice monitored over 6 months. Based on the DNA and RNA NGS global analysis of all the sequences in CRISPResso2, the most common mutation type is the single base deletion or insertion caused frame shifting in the coding sequence of *Myo7a* protein, which could result in disabling translation of *Myo7aSh1* proteins (Supplementary figures). The off-target analysis on the top 9 sequences showed very limited off-target effects. This work introduced a highly efficient and scalable EV loading platform for developing EV based precision gene editing with targeting progressive dominant genetic hearing loss as a disease model. The rapid loading of RNP complexes into EVs without using a dual vector could offer a great opportunity to customize sgRNAs addressing different mutant alleles within one gene, in turn, enabling customization to patients’ genetic heterogeneous mutational backgrounds, which holds clinical translational potential for overcoming current challenges in gene therapy.

## Results

### High throughput and highly efficient EV electro-transfection via the μDES platform

We developed the μDES platform for enhancing EV transfection efficiency, loading capacity, and throughput. The concept of the μDES platform is illustrated in Fig. 1a, which streamlines droplet generation as numerous bioreactors with continuous-flow electroporation controlled by a low voltage direct current (DC) power supply. Within the droplets, the cargo transportation into EVs mainly relies on the electrical mobility of the cargo themselves, electric flux, and concentration gradient. On such a small scale, mass transport for crossing transient pores in the EV membrane can be completed in milliseconds (Fig. 1b)^41,42^. The homogeneous electric distribution can be achieved within water-in-oil droplets which carries low conductive buffer as aqueous phase ^41,43,44^. As demonstrated by multi-physics COMSOL simulation in Fig. 1c-e, the electric field applies to each flow-through droplet for highly efficient electroporation. Uniform distribution of both flow and electric field enables the precision control over electro-transfection process. The device fabrication is facile and low cost using a 3D printer as detailed in the supplemental materials (Supplementary Figure 1). Continuous generation of uniform droplets at fast speed enables large-scale processing EVs for cargo loading, which has been demonstrated by collecting droplets in a large petri dish and 2mL Eppendorf tubes (Fig. 1f) in uniform droplet sizes (Fi. 1g). By using fluorescently tagged 100 nm polystyrene beads as the standard calibration over nano sized EVs, consistent and efficient encapsulation within droplets has been demonstrated in Fig. 1h. The low power electroporation did not alter the droplet size and quantity (Fig. 1i and j). For water-in-oil droplet generation, fluorinated oil FC40 and associated 2% w/w fluoroSurfactant were employed as the oil phase due to their low conductivity, chemical inertness and stability, and easy removal. Pharmaceutical grade FC40 oil is considered as the highly biocompatible and low-cost recipe for droplet generation in compliance with the FDA ^45,46^. We also introduced pharmaceutical grade trehalose as a stabilizer in the aqueous buffer during droplet electroporation to minimize EV aggregation, membrane fusion and leakage^24-28,47^. The droplet size which determines the throughput can be controlled by adjusting the pressure/flow rate of water-to-oil ratio^48^. For achieving high throughput, the droplets can be generated at high speed (∼700 droplets/min), which leads to ∼30mL per hour processing throughput for each device. Note that current cuvette electroporation only handles ∼100μL per sample. For efficient removal of excessive Cas9 cargo from loaded EVs, we employed Ni Sepharose high performance agarose beads to selectively capture the His-tagged Cas9 proteins in the suspension sample solution. The cargo loaded EVs in the aqueous phase can be harvested via low-speed centrifugation phase separation to fully remove the oil phase (Fig. 1a).

**Fig. 1.**
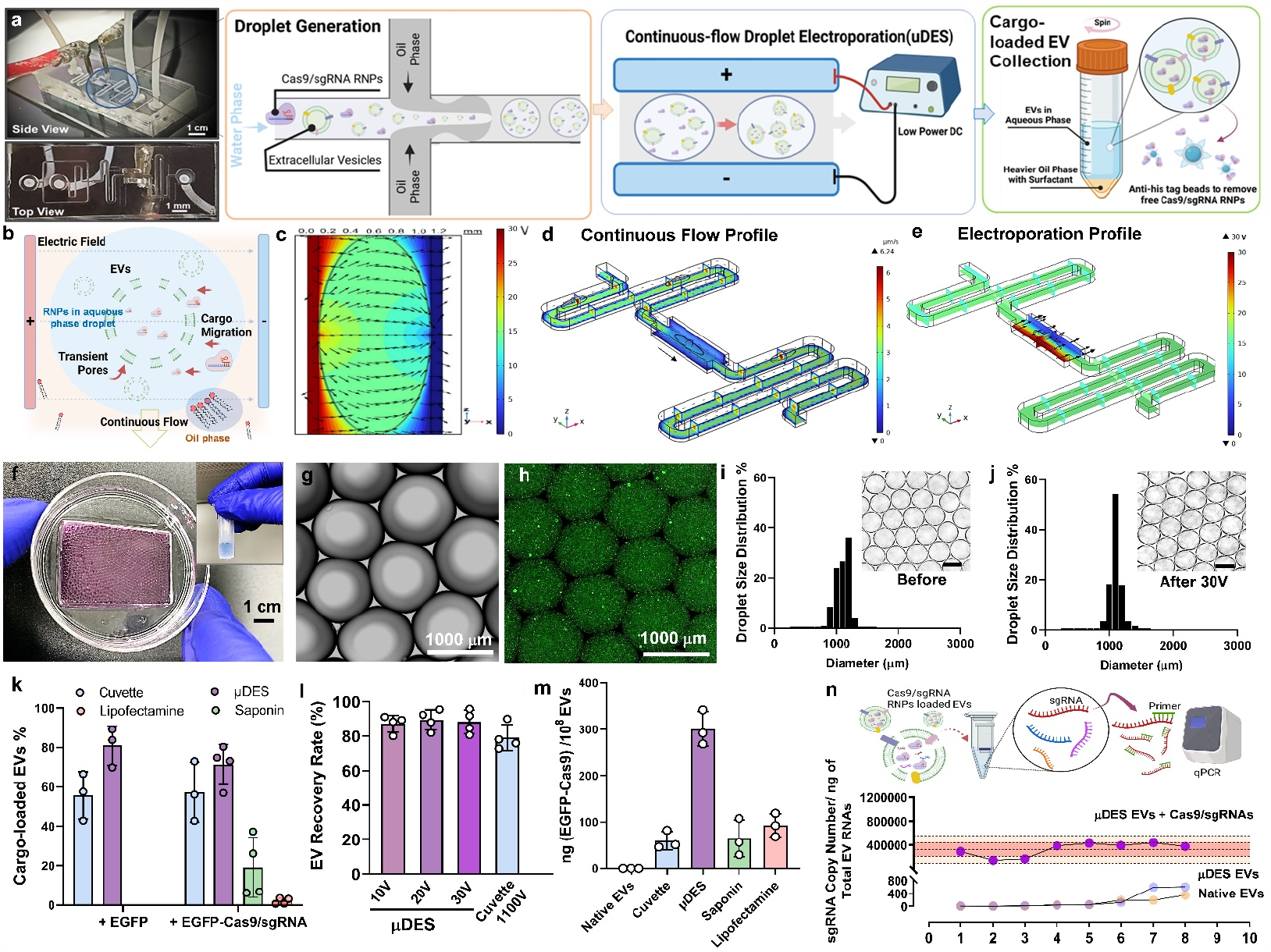
High throughput and highly efficient EV electro-transfection via μDES platform. **a**, Images of the μDES device with an illustration of continuous-flow droplet generation, droplet-based electroporation, and cargo-loaded EV harvesting and purification. **b**, Schematic illustration of droplet-based electroporation for EV cargo loading under uniform electric field distribution as demonstrated by multi-physics COMSOL simulation (**c**). **d**, The multi-physics COMSOL simulation analysis of continuous fluidic profile and electric field profile (**e**) to show the uniformity for precision control. **f**, Picture of large-scale collection of cargo loaded EVs from droplets. **g**, Microscopic image of continuous flow generated droplets with encapsulation of green fluorescence stained nanobeads (**h)** mimicking EVs. The droplet size is consistent and uniform before **(i)** and after (**j)** electric field application for transfection (30 V). The insert scale bar is 1000 μm. **k**, Evaluation of EV cargo loading rates among different transfection methods using fluorescence nanoparticle tracking analysis (fNTA). **l**, The recovery rate of μDES produced EVs compared with conventional cuvette electro-transfection which requires 1100V for electroporation. **m**, Quantitative measurements of Cas9 proteins from loaded RNP EVs normalized by EV particle number among different transfection methods. **n**, The quantitative PCR analysis of sgRNA copy number from loaded RNP EVs normalized by total EV RNAs. The electro-transfection was done by using the μDES platform in 8 replicates (Each has 3-4 technical replicates). The highlighted pink is 95% confidence region. The native EVs and μDES prepared EVs without RNP cargos both served as the negative control groups. EGFP-Cas9 were pre-assembled with gRNAs at a 1: 2 molar ratio before loading. EVs were purified from HEI-OC1 hair-like cell cultures for mixing with EGFP-gRNA:Cas9 RNP in electroporation low conductivity buffer. A final concentration of 10^10^ /mL of EVs was used for Neon cuvette electroporation control group, μDES system, and chemical transfection methods. Values and error bars represent the mean ± SD of three or more biological replicates.

For loading cargos into EVs, our platform achieved ∼80% rate for large proteins including gRNA:Cas9 RNP complexes, showing significantly higher loading efficiency than other conventional transfection methods including direct incubation, lipofection, and cuvette electro-transfection (Fig. 1k). The gRNA:Cas9 RNP cleavage bioactivity was not affected by μDES processing as assessed in Supplementary Figure 2 with the commercial cuvette Neon system as the control group. We also compared the recovery rate shown in Fig. 1l, which displayed a better recovery rate from μDES platform with only 30V voltage needed for electroporation. We quantified the EGFP-Cas9 protein loading amount (Fig. 1m) and compared it with conventional transfection methods, which showed more than 10-fold increase from μDES group. To characterize the reproducibility using μDES electro-transfection, we used well-established standard curve of chemically modified sgRNA (Supplementary Figure 2d) to quantify loaded gRNAs in the RNP-EVs via qPCR (Supplementary Figure 3). The native EVs and μDES conditioned EVs without RNPs served as control groups. The good loading capacity and reproducibility from 8 replicates were shown in Fig. 1n, without noticeable leakage or changes from intrinsic EV molecular content. Overall, the μDES platform demonstrated the advanced performance for EV cargo loading in terms of loading efficiency, throughput, and consistency.

### Characterization of μDES produced RNP EVs demonstrates high biocompatibility

The μDES platform maintained a consistent flow rate and uniform electroporation period for each droplet, which is essential for the reproducible and efficient cargo loading into EVs, in turn, retaining the natural EV properties as non-engineered EVs in terms of size (Fig. 2a, Supplementary Figure 4), zeta potential (Fig. 2b), protein contents (Fig. 2c), morphology and surface properties (Fig. 2d). Such results also indicate that μDES droplet oil phase does not impose noticeable influence on the EVs in the aqueous phase. We tested the essential protein content (CD81, TSG101, Alix) from μDES produced RNP EVs to compare with native EVs, which showed comparable expression level but carrying significantly high amount of encapsulated Cas9 proteins (Fig. 2c and Supplementary Figure 3d). The immuno-gold nanoparticle (AuNP) staining TEM imaging showed comparable morphology and surface expression of CD81 between RNP EVs and native EVs, without noticeable adsorption of Cas9/sgRNA RNPs on the EV surface, which indicates RNP cargoes were loaded inside of EVs (Fig. 2d). We also tested the biocompatibility (Fig. 2e) and cellular uptake behavior (Fig. 2f) from μDES produced RNP EVs sourced from both human bone marrow mesenchymal stem cells (RNP MSC-EVs) and HEI-OC1 hair cells (RNP HEI-OC1 EVs), compared with RNP LNPs group and native EVs. The μDES produced RNP EVs did not show significant differences from their un-loaded native EVs. Both EV groups (RNP MSC-EVs and RNP HEI-OC1 EVs) showed the ability to promote hair cell growth with superior cell biocompatibility compared with LNP groups. After one-hour cellular uptake, μDES produced EGFP-fused Cas9:sgRNA RNP MSC EVs exhibited higher uptake rate for cytoplasmic release and gradual entry into the nucleus (white arrow indication), compared with RNP LNP group (Fig. 2f and Supplementary Figure 5). Results suggest that EVs are advantageous for CRISPR RNP delivery with improved biocompatibility.

**Fig. 2.**
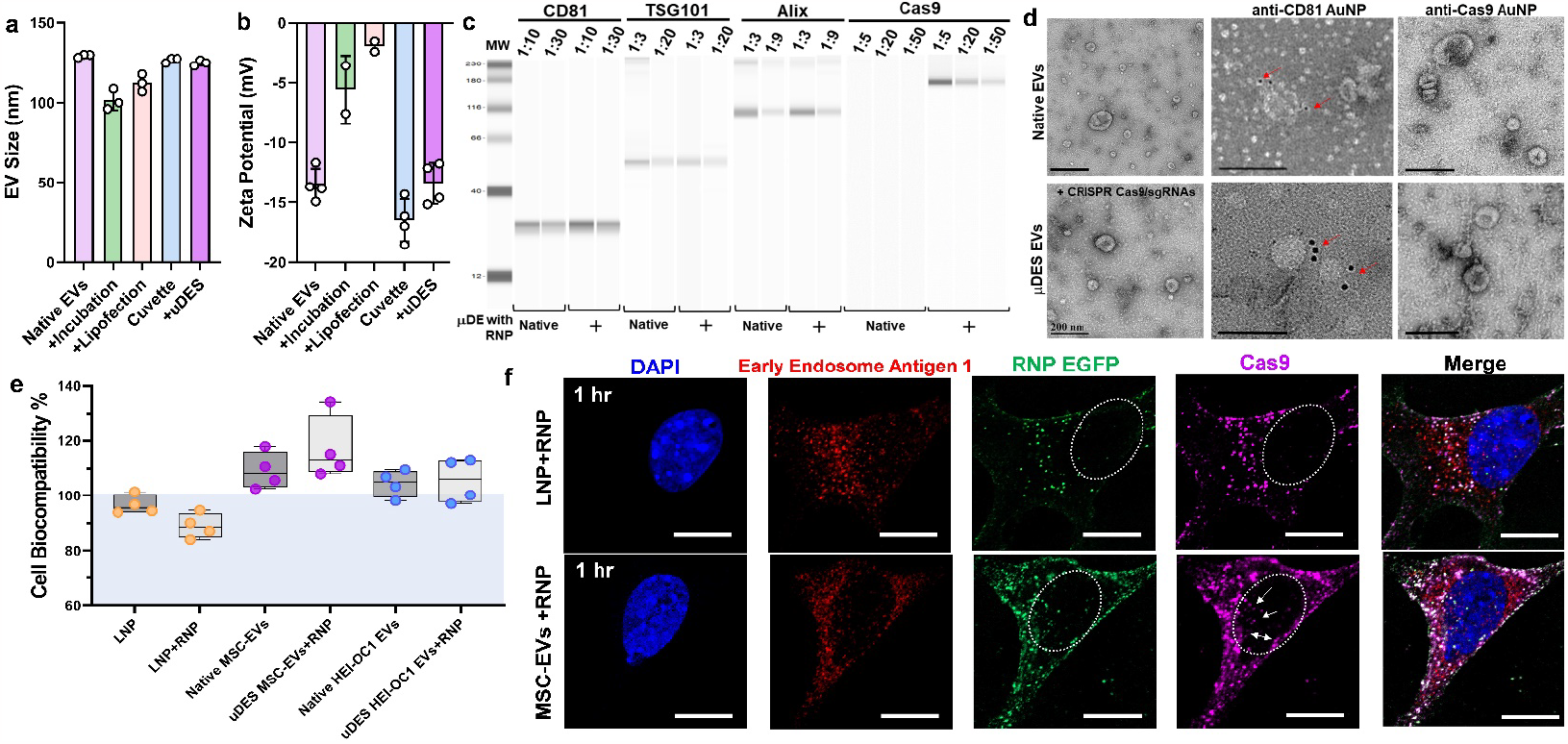
Characterization of μDES RNP EVs demonstrates high biocompatibility. **a**, Characterization of EV size and zeta potential **(b)** after transfection via different methods. The original native EVs serve as the control group. **c**, MicroWestern blotting analysis of essential EV protein contents (CD81, TSG101, Alix, Cas9) derived from μDES produced RNP HEI-OC1 EVs in serial dilution, with native EVs as the control group. **d**, Immune gold nanoparticle (AuNP) staining TEM imaging analysis of μDES produced RNP HEI-OC1 EVs with native EVs as the control group, in terms of CD81 surface marker expression and Cas9 surface adsorption. Scale bar = 200 nm. **e**, Biocompatibility analysis using PrestoBlue assay with HEI-OC1 hair-like cells (∼10^6^) dosed with LNP group (LNP-102, Cayman Chemical, w/o RNP), MSC-EV group (w/o RNP), and HEI-OC1 EV group (w/o RNP) in ∼10^9^ particles. **f**, Confocal imaging analysis of HEI-OC1 hair cells for uptaking particles in one-hour following dosing with RNP*^EGFP^ LNPs (LNP-102, Cayman Chemical) and RNP*^EGFP^ MSC-EVs in ∼10^9^ particles. The white arrow indicates the cytoplasmic release and gradual entry into the nucleus. Scale bar = 10 μm.

### CRISPR design for allele-specific editing of pathologic *Myo7a*^*sh1*^

*Myo7a* mutations make up 4.5% of cases of sensorineural hearing loss evaluated in a large human patient cohort^49^, which presents significant clinical populations with a high level of burden that is not addressable by current therapeutic interventions. Timely removal of mutant *Myo7a* allele could potentially prevent the progression of hearing loss. Therefore, we demonstrated an allele-specific editing system using gRNA:Cas9 RNP with designed gRNAs targeting G-C mutations in the Shaker-1 hearing loss mouse model. For *in vitro* validation, we used primary fibroblast cells isolated from *Myo7a* ^*WT/Sh1*^ mice and transfected either by electroporation of RNPs directly into cells (Fig. 3 and Supplementary Figure 6) or by internalization of μDES produced RNP EVs (Supplementary Figure 7). We screened Cas9 and two different gRNA sets harboring *sh1* mutation modified with 2’-O-Methyl and 3’-phosphorothionate bonds on the last bases on 5’ and 3’ end^50^ (Fig. 3a and Supplementary Figure 2d). In each gRNA set, we designed full-length and truncated forms of *Myo7a*^*sh1*^ targeted gRNAs (Table s1) to form the RNP complex individually for electro-transfecting primary fibroblast cells (Supplementary Figure 6a). The *in vitro* gene editing results demonstrated the efficient cleavage in the primary fibroblast cells with *Myo7a* ^*WT/sh1*^ and *Myo7a* ^*Sh1/Sh1*^ on the targeted *Myo7a* amplicons in the T7 endonuclease 1 assay (Supplementary Figure 6b), while displaying non-detectable indels in *Myo7a* ^*WT/WT*^. The indel percentage of gRNA-1 is reduced to ∼25% in *Myo7a* ^*WT/Sh1*^ fibroblast cells, compared to ∼45% indel percentage in *Myo7a* ^*Sh1/Sh1*^, which suggested robust editing selectivity from designed gRNAs when targeting *Myo7a*^*sh1*^ allele *in vitro*. Sanger and NGS sequencing were further performed to confirm the allelic cleavage specificity and editing efficiency (Supplementary Figure 6). The sequence percentage analyzed by CRISPResso2^50^ showed that gRNA-1 and gRNA-2 have higher cleavage activity than their truncated versions, respectively (Fig. 3b). The indel profile revealed that the majority of CRISPR-induced variants were deletions for gRNA-1 while insertions for gRNA-2 (Fig. 3c and e). The sequencing variations demonstrated that 94.83% of *Myo7a*^*WT*^ is unedited in *Myo7a* ^*WT/ sh1*^. In contrast, *Myo7a*^*sh1*^ sequences were greatly reduced to ∼18% and the edited sequences account for the rest 82% in the *Myo7a*^*sh1*^ allele. We then quantified the targeting specificity of gRNA designs by sorting out mutation pattern 5’-CCG-3’ and wild type pattern 5’-CGG-3’ separately. By designing PAM sequence of gRNAs which have closer proximity to the *Myo7a* mutation, ∼95% specificity was achieved for *in vitro* editing (Fig. 3d) and the wild-type allele is mostly intact after the gRNA:Cas9 editing (Fig. 3f). Based on the global analysis of all the sequences in CRISPResso2 (Fig. 3e and f, Supplementary Figure 7), the most common mutation type is either a single base deletion or a single base insertion causing a frame shift in the coding sequence of *Myo7a* protein, which could result in disabling translation of the *Myo7a*^*sh1*^ proteins^51^. The off-target analysis of the top 9 off targets demonstrated a very limited off target rate (Supplementary Figure 6e). For *in vitro* editing mediated by EVs and delivering RNPs into primary fibroblast cells, ∼6% editing percentage on *Sh1* allele was achieved using μDES produced RNP MSC-EVS, which is significantly higher than that from conventional cuvette electro-transfected RNP MSC-EVs (∼0.1%, Supplementary Figure 7f). Both transfections either through direct electroporation of RNPs or EV internalization resulted in similar levels of frame shifting in the coding sequence of *Myo7a* protein, indicating the stable gene editing behavior (Supplementary Figure 7e). Our EV mediated *in vitro* editing efficiency is slightly higher compared to the LNP-mediated editing efficiency in the fibroblast cells reported by Chen *et al* for mutant Tmc1 allele (∼4%)^52,53^. Taken together, full-length gRNA-1 and gRNA-2 have the best allele specific editing property, thereby, prompting further *in vivo* validation.

**Fig. 3.**
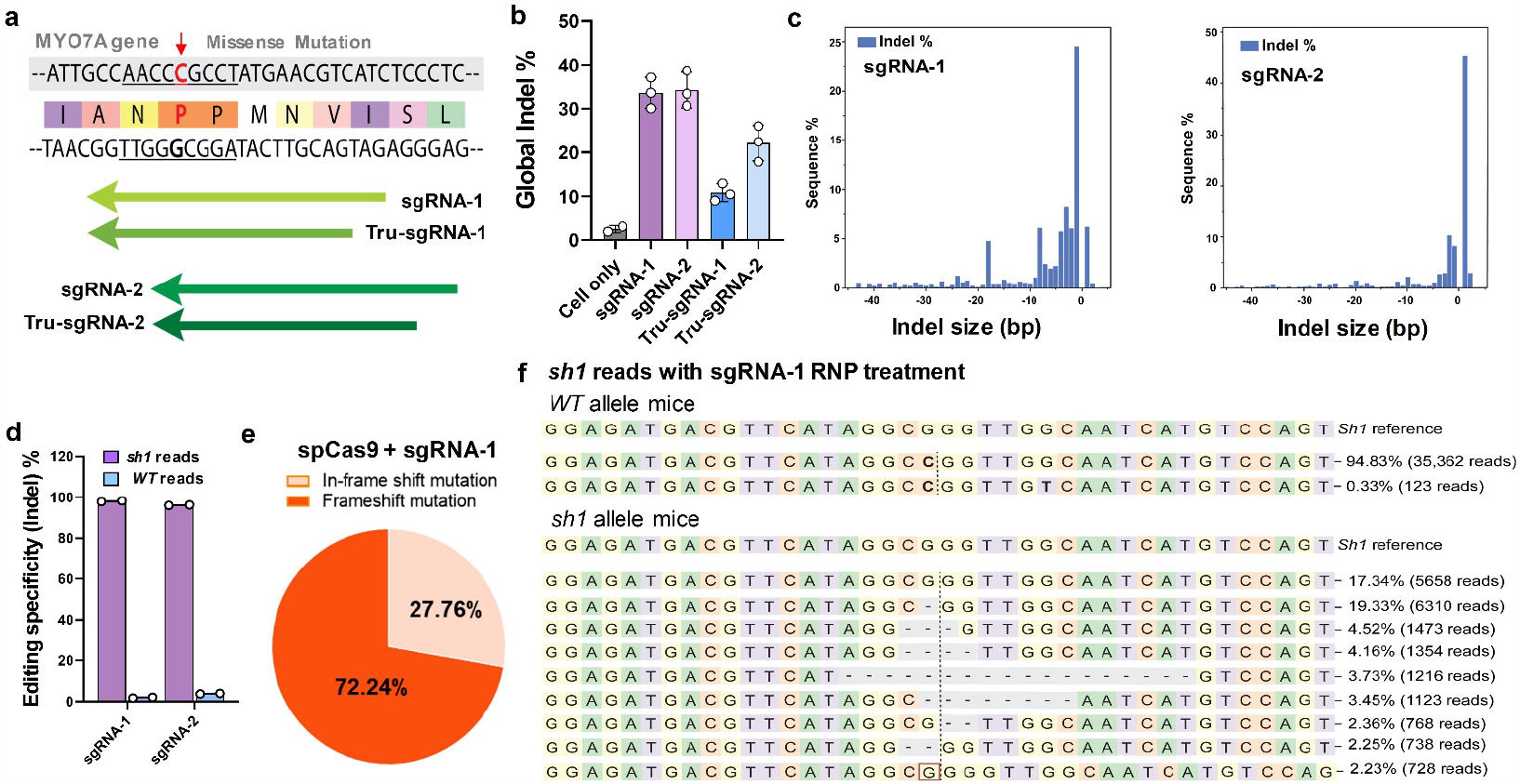
Design and characterization of CRISPR system for allele-specific editing on pathologic *Myo7a*^*sh1*^. **a**, Two different sgRNAs and their truncated version were designed to target the allele harboring Shaker-1 mutation. **b**, NGS results showed that editing indel% of gRNA-1 and gRNA-2 are ∼35%, which is higher than their truncated versions. **c**, Distribution of indel size revealed that gRNA-1 has more single-base deletions (-1) while gRNA-2 has more single-base insertion (+1). **d**, Allele specific analysis demonstrated that gRNA-1 and gRNA-2 displayed higher editing specificity to edit Shaker-1 allele. **e**, Pie chart showed the majority of editing events from sgRNA-1 are frameshift mutations with some in-frame mutations. **f**, Allele frequency tables demonstrated that shaker-1 allele is greatly reduced while WT allele is intact. All graphs show the mean ± SEM and individual biological replicates.

### μDES loaded RNP EVs for *in vivo* arrest of progressive hearing loss

The Shaker-1 mouse wild type *Myo7a* ^*WT/WT*^ and heterozygotes *Myo7a* ^*WT/sh1*^ initially both have normal hearing. However, the mutant heterozygotes *Myo7a* ^*WT/sh1*^ animals will gradually lose hearing ability with significant impairment testable at 6 months of age^37^. Thereby, Shaker-1 heterozygotes *Myo7a* ^*WT/sh1*^ serve as a good model for validating gene therapy *in vivo* in treating progressive hearing loss. In order to characterize the EV tissue biodistribution behavior in the inner ear, three groups of RNP*^EGFP^ LNP, μDES produced RNP*^EGFP^ MSC EVs and RNP*^EGFP^ HEI-OC1 EVs were injected via posterior semicircular canal, which delivers into the perilymphatic space of the inner ear (Fig. 4a). The μDES produced RNP-EVs displayed superior biocompatibility and rapid (within 2 hours) tissue delivery, particularly to the targeted outer and inner hair cells (Fig. 4b). In contrast, RNP-LNPs exhibited a tissue distribution limited to auditory nerve bundles and no delivery to inner and outer hair cells (Fig. 4b left). Thus, the results support the superior ability of μDES loaded CRISPR RNP EVs to target sensory cells in the inner ear. As illustrated in Fig. 4c, we administrated μDES produced RNP EVs (∼ 10^8^ particles in ∼1 μL) via the posterior semicircular canal into the left ear of *Myo7a* ^*WT/sh1*^ mice, with normal hearing *Myo7a* ^*WT/WT*^ mice and non-treated *Myo7a* ^*WT/sh1*^ mice as the control groups. Four weeks after EV delivery, the organ of Corti was extracted for Sanger, mRNA sequencing, and qPCR validation (Fig. 4d). NGS mRNA sequencing showed higher global indel% from both μDES produced RNP EV groups (MSC EVs and HEI-OC1 EVs) than that from RNP LNPs group (Fig. 4e), which indicates the improved gene editing ability *in vivo* using EV mediated gene delivery. For validation using qPCR to specifically measure the expression of *Myo7a Sh1* allele mRNA, both RNP EV groups lead to a significant reduction in treated mice compared to that from RNP LNPs group or untreated ears which showed no changes in gene expression (Fig. 4f). Such gene editing performance differences from EV and LNP groups could potentially be due to the tissue penetration ability of the particular LNP used for delivery. The editing efficiency was amplified at the mRNA level given the fact that mRNA of *Myo7a* was only transcribed in hair cells, although the *Myo7a* gene exists in every cell type in the organ of Corti ^54,55^.

**Fig. 4.**
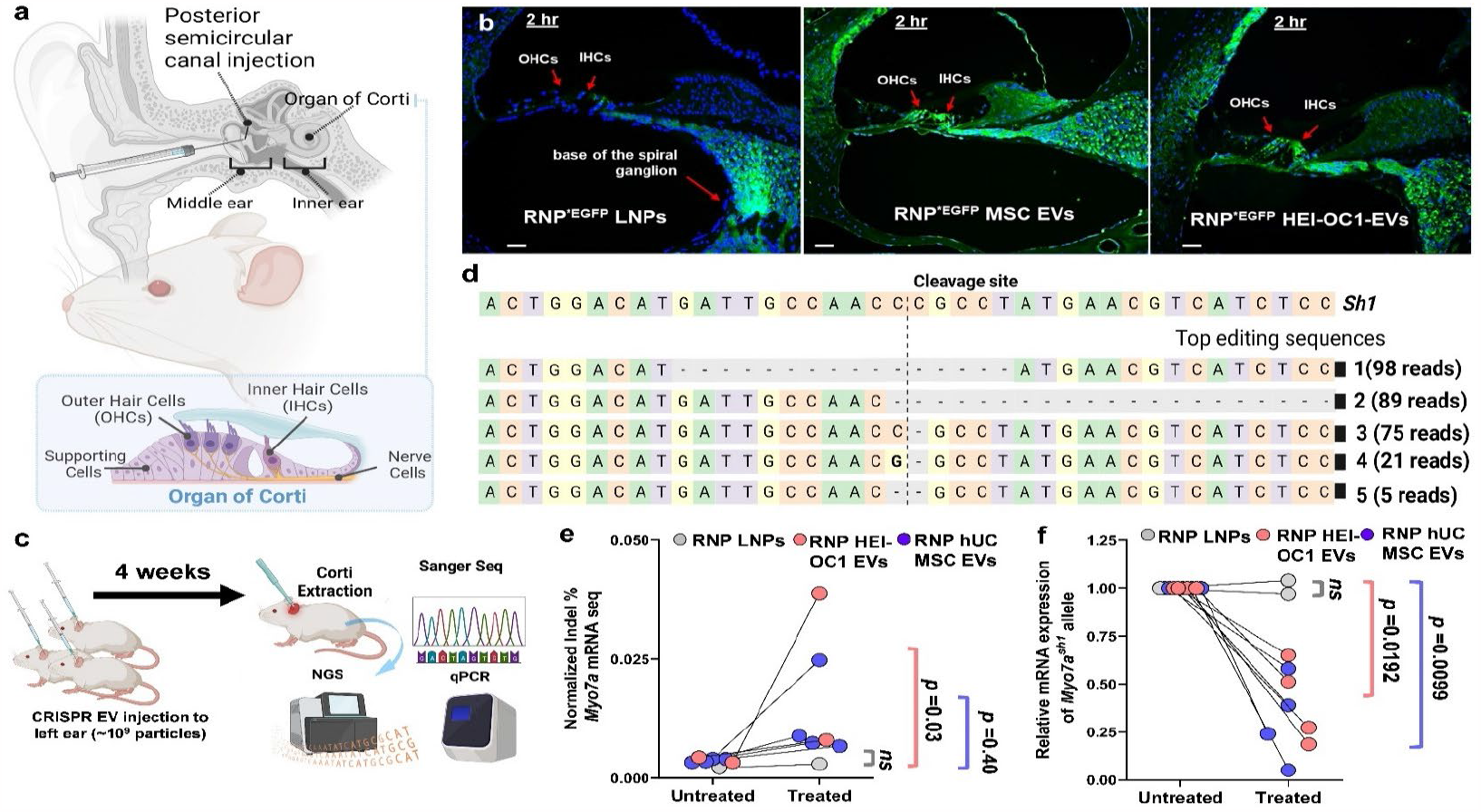
*In vivo* characterization of μDES loaded RNP EVs. **a**, Schematic illustration of ear structure and essential sensory outer hair cells and inner hair cells in the organ of Corti. The body of the hair cells are within the perilymph space whereas the stereocilia bearing apex of the cell protrude into the endolymph containing scala media. **b**, Confocal ear tissue imaging analysis of the biodistribution in the organ of Corti via posterior semicircular canal injection of RNP^*EGFP^ LNP (left panel), μDES produced RNP^*EGFP^ MSC EVs (middle panel), and RNP^*EGFP^ HEI-OC1 EVs (right panel) into Shaker-1 mice ears individually. Scale bar = 30 μm. **c**, schematic illustration of workflow for characterizing *in vivo* gene editing efficiency. **d**, Allele frequency tables demonstrated that shaker-1 allele is greatly reduced with top 5 editing sequences, while WT allele is intact. **e**, The global indel% from NGS mRNA sequencing on mice ear treated from μDES produced RNP MSC EVs and RNP HEI-OC1 EVs compared with RNP LNPs. The untreated mice ears served as the baseline control group. **f**, qPCR validation on the expression of *Myo7a Sh1* allene treated from μDES produced RNP MSC EVs and RNP HEI-OC1 EVs compared with RNP LNPs. The untreated mice ears served as the baseline control group.

We further monitored hearing ability after 6 months treated by single dose injection (∼ 10^8^ particles in ∼1μL) via the posterior semicircular canal into the left ear of *Myo7a* ^*Sh1/WT*^ mice and kept the right ear as the untreated control group. The auditory brainstem response (ABR) was measured with threshold (SPL) defined that at least one of the waves could be identified in 2 or more repetitions of the recording (Fig. 5b). For both treatment groups (RNP HEI-OC1 EVs in Fig. 5c and RNP MSC EVs in Fig. 5d), within the wide range of sound frequency from 4 kHz to 32 kHz, the treated left ears displayed significantly improved hearing threshold compared to the untreated right ears (∼ 20dB reduction in hearing threshold), which is nearly comparable to the healthy control group with the normal hearing ability (Fig. 5). The RNP MSC EVs treatment group (Fig. 5d) displayed slightly higher improvement than that from RNP HEI-OC1 EVs group (Fig. 5c), in terms of hearing restoration compared to untreated right ears (*p*=0.0009).

**Fig. 5.**
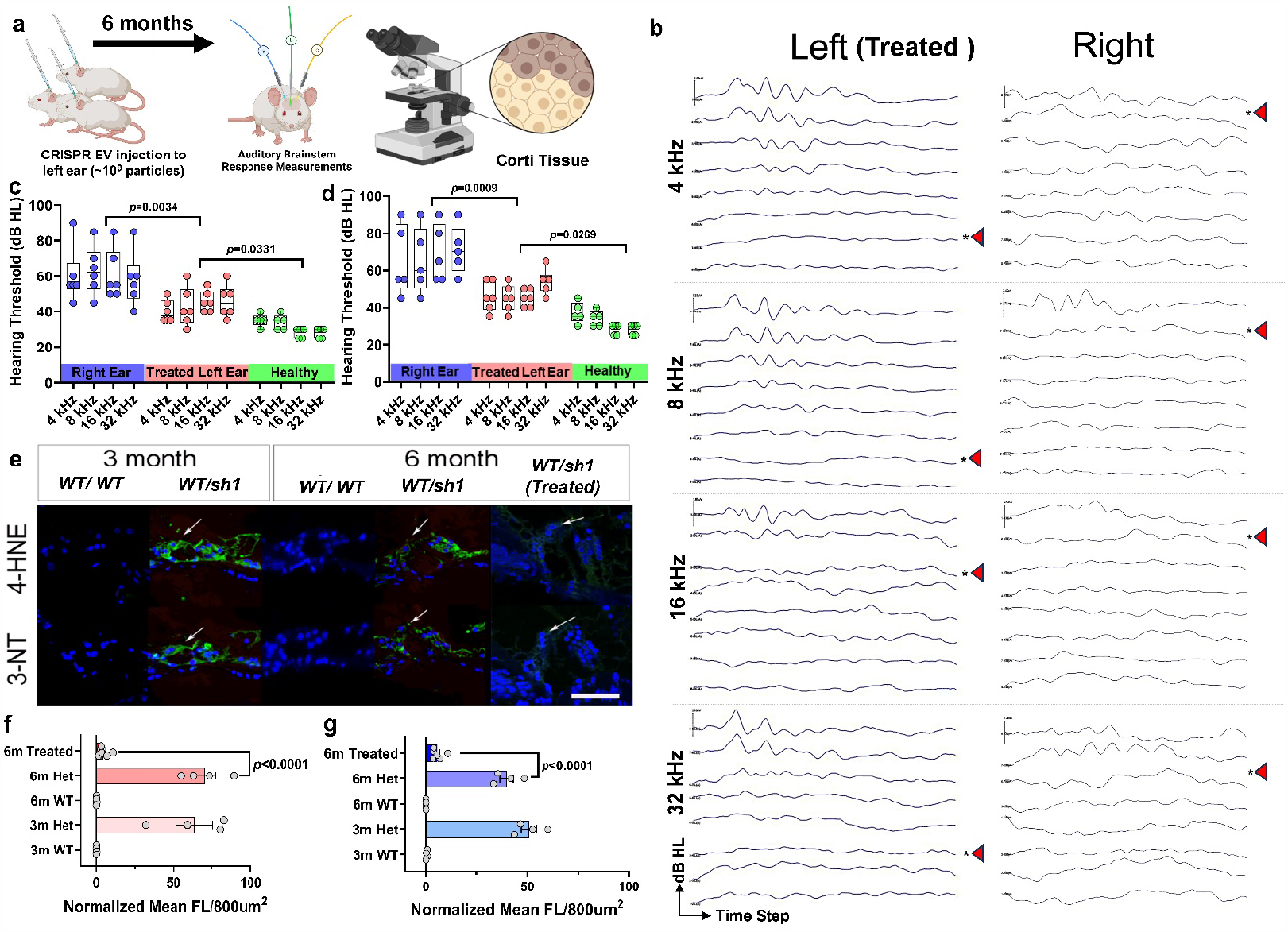
*In vivo* arrest of progressive hearing loss. **a**, Schematic illustration of animal auditory testing methods. **b**, Representative auditory brainstem responses (ABRs) recorded from shaker-1 heterozygotes mice left ear treated with μDES produced RNP MSC EVs in month 6, and right ear without treatment. The red arrow showed the hearing threshold. **c**, Groups of shaker-1 heterozygotes mice (n=6) in age of week 3 were treated with RNP HEI-OC1 EVs in left ear with right ear as the untreated counterpart, and wide type normal hearing mice as the control group. **d**, Groups of shaker-1 heterozygotes mice (n=6) in age of week 30 were treated with RNP MSC EVs in left ear with right ear as the untreated counterpart, and wide type normal hearing mice as the control group. The single dose injection (∼ 10^8^ particles in ∼1 μL) was administrated via the posterior semicircular canal. The ABR hearing was monitored 6 months after injection from sound frequency 4 kHz to 32 kHz. **e**, Confocal images from immunohistochemistry oxidative stress analysis of markers 4-HNE and 3-NT stained in the inner ear hair cells in Corti, at 3 and 6 months of age mice (n=6) from both hearing disable heterozygotes (*WT/ sh1*) and normal hearing wide type (*WT*/*WT*). **f**, Quantitative analysis of expression level of oxidative stress markers 3-NT and 4-HNE **(g)** in the inner ear hair cells from Corti, at 3 and 6 months of age mice (n=6) from both hearing disable heterozygotes (*WT/ sh1*) and normal hearing wide type (*WT*/*WT*).

Our previous work has shown a correlation between cochlear oxidative stress and genetic hearing loss ^56,57^. Therefore, we evaluated the Shaker-1 mouse inner ear for the presence of the oxidative stress markers 4-Hydroxynonenal (4-HNE) and 3-Nitrotyrosine (3-NT). The heterozygous mice (*Myo7a* ^*WT/ sh/*^) showed oxidative stress marker labeling throughout the organ of Corti and higher expression levels of both oxidative stress markers at 3 and 6 months of age in the inner ear hair cells (Fig. 5 e-g, white arrows), in contrast to normal hearing wild type (*WT/WT*) mice. In our treatment group of heterozygous (*WT/sh1*) mice, both 4-HNE and 3-NT expression levels were significantly reduced to unnoticeable level at 6 months of age (*p*<0.0001), indicating minimized oxidative stress and associated improvement of hair cell function after gene editing. Overall, the result supports that EV mediated CRISPR delivery and in vivo editing is an advanced and therapeutically effective approach.

## Discussion

Development of treatments for dominant hearing loss has significantly lagged behind compared to treating recessive disorders. Recently, genome editing brought a promising approach to correct genetic defects in dominant hearing loss using CRISPR Cas9 technology. However, for clinical translation, a major hurdle is the successful packaging of these larger endonuclease systems for effective *in vivo* delivery. Presently, significant challenges with the application of viral vector or LNP-based delivery into the auditory sensory system have limited their rapid clinical translation in treating hearing loss. Using EVs to encapsulate gRNA:Cas9 gene editing agents provides a transient way to target the inner ear for achieving permanent genetic alterations of deafness genes. EVs have excellent tissue penetration ability, outperforming LNPs as reported previously^58^. We also observed the specific biodistribution of EVs in the inner ear tissue for accessing and targeting inner ear cells as opposed to LNPs in Shaker-1 mouse model (Fig. 4). EVs also targeted other inner ear tissues such as the stria vascularis and the spiral ganglion. By investigating the therapeutic potential of RNP EVs, we have demonstrated the *Myo7a* ^*WT/ Sh1*^ editing ability from RNP EVs in the progressive hearing loss mouse model. Combined with *in vivo* hearing recovery over 6-month monitoring, the results showed that RNP EVs can disrupt the expression of *Myo7a*^*sh1*^ allele at the mRNA level, thereby attenuating the oxidative stress that is associated with progressive hearing impairment. We observed the reduction of oxidative stress markers in both inner and outer hair cells of RNP EV treated cochlea (Fig 5), suggesting effective editing and reduction of Shaker-1 allele induced pathology in both cell types. The ability to deliver into a clinically accessible space in adult animals and accessing both inner and outer hair cells highlights the translational potential of this approach. Most importantly, the treatment of RNP EVs on the *Myo7a* ^*WT/ Sh1*^ mice also exhibited a promising reduction in the hearing threshold. Potentially, RNP EVs could be applied to a range of different genetic defects associated with hearing loss. This novel delivery strategy avoids potential ototoxicity induced by delivery vehicles or overexpression of gRNA:Cas9 complexes caused off-target effects *in vivo*.

Allele-specific gene editing via CRISPR-Cas9 was recently shown to successfully prevent hearing loss by disrupting the mutated transmembrane channel-like gene family (*TMC1*) allele in *Beethoven* mice using a variety of delivery approaches including Cas9-guide RNA-lipid complexes^59^, AAV vectors^60^, and RNP LNPs^52^. The neonatal mouse is often used in those studies which could vary in terms of delivery distribution when comparing with the adult mice, as the hearing matures after postnatal day 14. Using RNP EVs, we demonstrated successful prevention of hearing loss phenotype in both neonatal (Fig. 5c) and adult animals (Fig. 5d). Additionally, we utilized a delivery approach (injection into the perilymph space) that can be translated into human patients clinically. Based on human cochlear implantation data, in a mature ear, violation of the scala media (which has been used in other delivery experiments) results in loss of residual function, making delivery to the perilymph space essential^61^. Interestingly, we were unable to achieve any significant delivery of RNP LNPs to the inner ear hair cells using a perilymph-based delivery (Figs 4 and 5). Using our delivery approach, LNPs were limited to the basal region of the spiral ganglion (Fig 4). As shown by others ^62^, EVs derived from different organisms can cross-traffic between kingdoms between plants, microbes, animals and humans. We tested mouse HEI-OC1 cells derived EVs and human mesenchymal stromal cells derived EVs *in vivo* and both showed effective distribution to a wide range of cells within the inner ear including inner and outer hair cells (Fig 4), which opens more options of cell sources for EV production.

Additionally, to understand the potential impact of *in vivo* gene editing with in-frame mutations on protein structures, we employed Alphafold2 pipeline with relaxation to characterize the structural similarity to the residue 401-561 in the wild-type motor domain where harbored shaker-1 mutation along with Cas-induced mutants in Supplementary Figure 8 and 9^63^. Jointly considering the loop region was mostly tolerant to the indel in general^64,65^ and the basis of AlphaFold2’s prediction^63,64,66^, we expect that 1-residue deletion on the loop region is unlikely to reduce significant structural integrity compared to WT domain shown in the Supplementary Figure 8. Similarly, the pLDDT of 6-residue deletion of the loop region also showed high tolerability to the deletions and high foldability, while the binary alignment to WT exhibited slightly different positioning 6-residue deletion domain in Supplementary Figure 8. By reducing percentage of Shaker-1 allele from Cas9-mutagenesis, WT *Myo7a* allele to Shaker-1 allele is notably enhanced both *in vitro* and *in vivo*. Therefore, the functional WT *Myo7a* proteins can dominate over the possible variants if there is any. Such domination is greatly beneficial for the prevention of hearing impairment. Though it is still challenging to predict the gradual or drastic shifts on the function of final in-frame mutants based on the computational toolkits,^64^ the residual expression level and the associated high structural integrity render potentially limited impacts of in-frame mutants on the biological function of hair cells for hearing. Such results support the safety of our *in vivo* gene editing strategy.

Successful utilization of EVs for clinical translation as therapeutics highly relies on the capacity of cargo loading and production throughput. Our developed μDES platform has demonstrated 10-fold enhancement of loading efficiency and more than 1000-fold increase in processing throughput on loading RNP complexes into EVs, compared with conventional transfection methods. EVs’ membrane consists of a lipid bilayer with similar membrane protein composition as their donor cells. However, nanosized EVs contain more compact membrane curvatures and strong Brownian motion. Using microfluidic droplet-based electroporation could fully take advantages of efficient mass transport in confined space to maximize loading capacity, in turn, deriving optimal gene editing efficiency *in vitro* and *in vivo*, as we demonstrated in Fig. 4 and Fig.5. Fast continuous flow through of droplets also prevents direct contact of EVs with electrodes and avoids any thermal damage on EVs to retain EV natural integrity and stability. In contrast to kilovolt high voltage in conventional workflow, our technology substantially minimizes Joule heating and high voltage risk with using only 10-30 volts. The uniform electric field can be precisely controlled in compact droplet space, which can greatly improve consistency and transfection efficiency. Such a platform offers an easily amenable approach for scaling up by integrating multiple chip units for future GMP grade manufacturing of various cargo loaded EVs. Our approach will set as the first enabling gene therapy for tailoring mutation targets and advancing personalized precision gene therapy.

## Methods

### Materials and reagents

All chemical reagents and materials were purchased from ThermoFisher unless otherwise specified. Modified gRNAs were synthesized and quantified by Synthego. NLS-Cas9-NLS (Z03469), EGFP-Cas9-NLS nucleases (Z03467), and anti-Cas9 antibody (Clone 4A1) were purchased from Genscript. All DNA oligos were purchased from Integrated DNA Technologies. Collagenase IV was obtained from STEMCELL Technologies. SYLGARD™ 184 Silicone Elastomer Kit was purchased from Dow Silicones Corp. Master Mold for PDMS device was obtained from CADworks3D. Q5 high-fidelity DNA polymerase was purchased from New England Biolabs. Human bone marrow-derived conditioned culture medium and human umbilical cord-derived conditioned culture medium for extracellular extraction were purchased from EriVan Bio. SM-102 LNP in ethanol was a generous gift from Dr. Fan Zhang, in the College of Pharmacy at the University of Florida. 6nm Goat anti-Mouse and anti-Rabbit, IgG, Immuno Gold reagents were purchased from AURION.

### CRISPR Design and gRNA synthesis

CRISPR gRNA sequences were designed by using the Custom Alt-R CRISPR-Cas9 guide RNA provided by IDT (https://www.idtdna.com/site/order/designtool/index/CRISPR_SEQUENCE) and Horizon CRISPR Design Tool (https://horizondiscovery.com/en/ordering-and-calculation-tools/crispr-design-tool). Two gRNAs that harbor the single mutation in *Myo7a* gene were selected, and the truncated version of the corresponding gRNAs that is 3nt shorter were also investigated accordingly^67^ shown in Table 1. The sequences were then synthesized by Synthego Inc. with the following modifications, 2’-O-Methyl at three first and last bases, 3’ phosphorothioate bonds between first three and last two bases^50^. gRNAs were then dissolved in nuclease-free TE buffer (10mM Tris, 1mM EDTA, pH 8.0) to make the final concentration of 100 μM and stored in -20°C for future use.

### Cell culture and transfection

Mouse primary fibroblast cells were isolated from ear punches from C57BL/6^*WT/WT*^ (*Myo7a*^*WT/WT*^), heterozygous Shaker-1^*sh1/WT*^ (*Myo7a*^*sh1/WT*^) and Shaker-1 ^*sh1/sh1*^ (*Myo7a*^*sh1/sh1*^). After euthanasia, a small piece of external ear was cut out and sterilized with 70% ethanol for 5 min in a sterile 15 mL conical tube. Small ear punches were then washed with Hanks’ Balanced Salt Solution (HBSS, ATCC) without Ca^2+^, Mg^2+^ and then dissected into smaller pieces with razor. Next, mashed tissue suspension was further treated with ∼270U of prewarmed collagenase IV (STEMCELL Technologies, USA) in a cryotube vial for 90 min at 37°C followed by 0.05% trypsin-EDTA for 20 min at 37°C. Isolated fibroblast cells after filtered through a 70-μm mesh filter were then cultured in 10% fetal bovine serum containing DMEM (Gibco, USA) and supplemented with 1% GlutaMax (Thermo, USA) with 5% CO_2_ at 37°C. Fibroblast cells were validated with Sanger Sequencing (Genewiz, USA) before electro-transfection. For electro-transfection, Neon platform (Thermo, USA) was optimized and used for loading gRNA:Cas9 into the fibroblast cells according to the manufacturer’s instruction. Briefly, 60K fibroblast cells detached from the flask with 0.25% trypsin-EDTA were resuspended in R buffer in 10μL Neon electroporation kit. The preformed gRNA:Cas9 complexes were then added into electroporation solution to make 2 μM Cas9 and 6 μM gRNA in the final electroporation buffer. The electroporation was performed with 1500 V, 30 ms with 1 pulse and then cells were gently put into a 12-well plate containing prewarmed culture medium, incubated at 37°C, 5% CO_2_ for 96 hours.

### Preparation and isolation of extracellular vesicles

HEI-OC1 cells were subcultured in 10% fetal bovine serum supplemented DMEM (Gibco, USA) in T-175 (175 cm^2^) flasks at 33°C, 10% CO_2_ to reach 60% confluence. The culture medium was then replaced with 10% exosome-depleted fetal bovine serum containing DMEM (Gibco, USA) for EV production after washing cells twice with cold 1x PBS buffer. After 48 hours, the cell culture medium was collected in 50mL sterile conical tubes and then centrifuged at 350 xg, 2000 xg, 10,000 xg for 5min, 20min, 30min respectively, to remove cells, cell debris and microvesicles. The resulting medium was then stored in -80°C or further ultracentrifuge. To isolate certain size range of extracellular vesicles, 35% (w/w) sucrose in cold 1x PBS buffer without Ca^2+^, Mg^2+^ was prepared freshly and sonicated for 5min to fully dissolve sucrose pellet. The resulting sucrose solution was then filtered with 0.2μm PVDF membrane filter before use. 3mL of 40% sucrose solution was added to form the sucrose bed beneath 15mL cell culture medium in polycarbonate thick-walled tubes (Thermo, USA). After 90min ultracentrifuge at 100,000 xg, 3mL sucrose bed was transferred to a new ultracentrifuge tube and then disrupted in another 15mL cold 1x PBS buffer for another 90min ultracentrifuge. The resulting EV pellets were resuspended in 1mL cold 1x PBS buffer and filtered with 0.2 μm PVDF membrane filter before quantification and size distribution analysis with ZetaView® nanoparticle tracking analysis system. All centrifugation steps above were conducted at 4°C.

### Droplet generator on chip

The 3D structure of microfluidic device was designed and drawn by SOLIDWORK CAD. The resin mold for PDMS casting containing the feature of 1mm electrode channels, 150 μm in width as nozzle for droplet generation, was printed by μMicrofluidics Printer (CADworks3D, 30 μm resolution) shown in Figure S1. Briefly, CAD files was opened by *utility*.*exe* that is connected to μMicrofluidics Printer. The software setting for printing as follows: 50 μm as thickness, 0.1 for grid size, 40% Power ratio. The microstructure was then sliced and launched for printing. The resulting resin mold after the printing was then soaked in fresh 100% Ethanol or isopropanol for 20-30min 2-3 times to remove the free resin before the final UV curing step. And then the resin mold was dried with compressed Nitrogen. The soaking-drying cycles have to be repeated several times until there is no shiny free resin on the microstructure. Each side of resin mold was then cured in the Creative Cure Zone (CADworks3D) for another 10min twice for the final photopolymerization and solidification of microstructures. The resin mold was then ready for PDMS casting. PDMS was prepared using the standard 10:1 (base to curing agent) ratio. The PDMS mixture was stirred completely at least 3min and then degassed for at least 30min before being poured into the 3D-printed molds and then baked at 75°C for 3 hours. After the surface activation of molded PDMS pieces using a corona discharger and the microscope plate, PDMS molds were then assembled and bound onto the microscopic plate as the droplet-based electro-transfection device.

Electrodes which was made with 90% foot platinum and 10% iridium (SUREPURE CHEMICALS LLC, USA) were tailored to ***L*** shape to fit into electroporation chambers designed in μDES and then carefully inserted into the electroporation sites to ensure precise alignment. The 1/16 OD, 1/32 ID tubings were then inserted into the inlets and outlets in μDES. To avoid any potential leakage that can result in the unstable flow rate in μDES, the additional PDMS was then added to the area surrounding the inlets, outlets and electroporation sites before use. The device was then baked in the oven at 75°C for another 30min for the following electroporation.

### COMSOL Simulation

The proposed microfluidic device was fine-tuned using the COMSOL Multiphysics software package. The device’s mathematical model involves fluid flow and electromagnetism. For both models a standard linear triangular extra-fine mesh was assigned to the geometry. To observe the geometric evolution of our droplets, we used the computational fluid dynamics system (CFD) module using the laminar two-phase flow. For this simulation, the oil phase material was defined as FC-40 with a density of 1850 kg/m^3^ and a dynamic viscosity of 0.0018 Pa/s. Microfluidic flows are defined by the Navier Stokes equation where *ρ* is the density of the fluid, *u* is the velocity of the field, *t* is time and *P* is the pressure field:

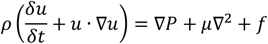

The physics of the electroporation system of the device were defined using the AC/DC module to simulate the electric field distribution in the microfluidics electroporation model. The material of the droplet was defined as electroporation buffer with a conductivity of 1x10^−4^ S/m. The electrical conductivity of the electrode was defined by the composition of the Platinum-iridium wire as 9.43x10^6^ S/m. The measurement of FC40 supplemented with 2% FluoroSurfactant was performed in the voltmeter and the conductivity of oil phase was approximated as 1.044 S/m. In steady conditions, the flow of electric currents within a conducting fluid follows Ohm’s law:

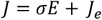

Here, *J* represents the total current density within the material while *σ* represents the electrical conductivity measured in S.m^-1^. *σ* is the electrical field strength measured in V.m^-1^ and *J*_*e*_ represents the current density.

Moreover, to mathematically describe the electric field that acts upon the droplet within the microfluidic device, we used the induced potential difference ΔΨ_*i*_ at a point *M* of the droplet membrane at any time *t*

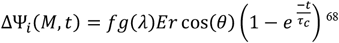

The results of this simulation were then used to adjust and optimize the device’s design for the intended application. Overall, the use of the COMSOL Multiphysics software package allowed for a detailed and accurate analysis of the proposed microfluidic device, ensuring its optimal performance.

### Bioactivity Assays of electroporated CRISPR/Cas9 RNP

The designed 6 μM CRISPR gRNAs were assembled with 2 μM Cas9 nucleases at room temperature in Neon R buffer for 10 min and then treated with electric filed with 1500 V, 30 ms and 1 pulse with 10 μL electroporation tip. The resulting CRISPR/Cas9 RNP were then transferred to 1.5mL Eppendorf tube and recovered for 10 min at room temperature. 200 ng of purified *Myo7a* amplicons were then added to 0.5 μM CRISPR/Cas9 RNP and then incubated at 37°C for 30 min. 2μg proteinase K (Thermo, USA) was then used to degrade Cas9 protein for 15 min at 37°C and followed by 2μg RNase A (Thermo, USA) to completely deactivate CRISPR/Cas9 RNP at 37°C. The final solutions were heated at 75°C for 10 min to end the enzymatic digestions. The final cleaved products were analyzed in 0.8-2% E-gel electrophoresis system (Thermo, USA) and imaged with Typhoon imager with Cy2 channel.

### Loading of RNP to EV

#### Neon electro-transfection system

EVs in 1x PBS buffer were firstly transferred into Neon R buffer (Thermo, USA) by using 30K cutoff ultrafiltration column to the final concentration of 10^10^ EVs/mL. Basically, EVs were added into the pre-washed 30K cutoff column and centrifuge at 7000 xg for 8min and then resuspend the concentrated EV (∼80μL) in 400μL Neon R buffer to centrifuge again under the same condition. The resulting EVs in Neon R buffer (∼60μL) was placed on ice immediately for later use. To electro-transfect CRISPR/Cas9 RNP into EVs, gRNAs and EGFP-Cas9 nuclease were firstly pre-mixed together in 45μL Neon R buffer and self-assembled at room temperature for 10min and then added into EVs solution to obtain 6μM of EGFP-Cas9 and 9μM of sgRNA in the ready-to-electro-transfection solution (∼106μL). To stabilize the membrane, 4.4μL of 1250mM trehalose in PBS buffer without Ca^2+^ and Mg^2+^ was added to the ready-to-electro-transfection solution to have 50mM trehalose in the final solution. The addition of trehalose increased the viscosity of electro-transfection solution therefore, to maintain the viscosity balance in Neon electroporation system, 120μL of 1250mM trehalose was added to 3mL Neon electrolytic buffer. 1500V, 20ms, 1 pulse was used to electro-transfect CRISPR/Cas9 RNP into HEI-OC1 derived EVs with 100uL Neon platform. The resulting RNP-EV was gently transferred to 1.5mL protein low binding Eppendorf tube for membrane recovery at room temperature for 10min. And then 500uL PBS buffer at room temperature was added to EV solutions followed by the further membrane recovery at 37°C for 20min. The resulting RNP-EV was stored in -20°C for the downstream purification and analysis.

#### High-throughput droplet-based μDES

Water-in-oil droplets were generated at the flow-focusing junction inside the μDES. As described previously, 10^10^/mL EVs were first transferred to Cytoporation® media T (Biochrom Ltd., United Kingdom) and then mixed with 5μM sgRNA-Cas9 RNP (gRNA:Cas9 molar ratio=1.5:1). The resulting mixture was delivered into the device as the dispersed aqueous phase. The oil phase contained FC-40 mixed with 2 weight % 008-FluoroSurfactant (RAN Biotech). A microfluidic pressure flow controller (PreciGenome LLC, USA) was used to generate the droplets with the diameter of around 1000μm at 3.0-3.8μL/min of the aqueous solution and 1.5-2μL/min of the oil phase. The electroporation on device was performed ranging from 10-60V by using direct current-based power supply (GW INSTEK, USA) and the resulting emulsion was collected within an Eppendorf tube or the microplate analyzed under the inverted microscope Cytation 5 (BioTek, USA). The isolation of aqueous phase containing RNP-EV from the oil phase was performed under the centrifuge at 2000-3000xg for 5-10min at room temperature. The aqueous phase was then collected by the pipettes and transferred to a new Eppendorf tube for downstream purification. The resulting RNP-EV was stored in -20°C for the downstream purification and analysis.

#### Chemical transfection methods for RNP-EV

The same amount of RNP mentioned above was firstly premixed and then coincubated with 2% saponin as reported before.^69^ For loading of RNP to EVs with lipofectamine 3000, 5μM RNP was mixed with 1μL lipofectamine 3000 and incubated for 20min at room temperature according to the manufacturer’s protocol for DNA plasmid transfection. The resulting RNP-EVs were then purified as described below.

#### Preparation of RNP-LNP

The same amount of RNP mentioned above was firstly premixed and then added to 1/10 volume of premixed SM102-LNP (Cayman Chemical, USA) according to the manufacturer’s instruction for the mRNA loading. For loading of RNP to EVs with lipofectamine 3000, 5μM RNP was mixed with 1μL lipofectamine 3000 and incubated for 20min at room temperature according to the manufacturer’s protocol for DNA plasmid transfection. The resulting RNP-EVs were then purified as described below.

### Purification of delivery vehicles RNP-EVs and RNP-LNP

To remove excessive His tagged CRISPR/Cas9 RNP from RNP-EVs and RNP-LNP, 100μL Ni Sepharose high performance beads (GE Healthcare) were firstly washed with 10 volume of cold 1x PBS buffer and then pre-equilibrated in PBS buffer for 10min. The equal volume of beads were then incubated with RNP-EVs or RNP-LNP at 4°C for 0.5-1 hours on the rocker until no more binding between His tagged EGFP or EGFP-CRISPR/Cas9. The unbound RNP-EV was then collected in 1mL of cold 1x PBS buffer while the excessive CRISPR/Cas9 bound on the column. The purified RNP-EVs and RNP-LNP were subsequently concentrated to a 2-fold increase using a 30K ultrafiltration column under the same conditions as described earlier and stored in -80°C for further analysis.

### Characterization of RNP-EV

#### Nanoparticle tracking analysis

The size and particle number of purified EV and RNP-EV were analyzed by nanoparticle tracking analysis (NTA) using the ZetaView® (PARTICLE METRIX, Germany) supplied with multilaser system. Briefly, to measure the size and number of EVs in the scatter mode, the EV solutions was diluted with 1x PBS to reach the meansuring concentration of 4-8x10^7^/mL via a blue laser (488nm). Subsequently, 1-2mL diluted EV solutions were then injected into the measuring cell and each measurement of every sample was repeated 2-3 times. Data analysis was carried out in ZetaView software with the following software settings for capture and analysis: Sensitivity=80, Shutter=100, Min brightness=20, Max brightness=1000. To quantify the EVs that were encapsulated with EGFP or EGFP-CRISPR/Cas9, fluorescent NTA analysis was performed by increasing sensitivity to 90 and switching to a 505nm filter while keeping the other parameters unchanged.

#### Zeta potential

The zeta potential of the purified RNP-EVs was measured by via electrophoretic light scattering (Litesizer 500, Anton Paar). Briefly, 25 μL of the final EV solutions diluted in 1:20 with 10% PBS buffer was injected into disposable folding capillary cuvette and each zeta potential measurement was conducted five times. The final measurement is conducted with 2min for pre-equilibrium, under room temperature by selecting PBS as the referenced conductivity.

#### Loading efficiency of Cas9 protein

The concentration of CRISPR/EGFP-Cas9 loaded into EVs was measured by Cytation5 (BioTek). Firstly, the standard curve of fluorescent intensity of EGFP-Cas9 in PBS buffer was obtained from Cytation 5 with a serial dilution of EGFP-Cas9 in the 96 well microplate. Each dilution was conducted and measured in duplicates. 35uL of the final EV was diluted in 1:1 with PBS buffer and then added to the microplate above and the fluorescent intensity of the resulting solution measured under the same condition with the standard curve. To further quantify the loading efficiency of CRISPR/EGFP-Cas9 in single EV, 25uL of the final EV solutions diluted in 1:40 with PBS buffer was then injected into ZetaView NTA (ParticleMetrix, Germany) for the quantification of EGFP^+^ EV. Basically, the diluted EVs were measured in scattering and fluorescent mode by using particle number/sensitivity measurement in fluorescent NTA (fNTA) with laser 488nm. By using reference polystyrene beads with Ex 488nm, the sensitivity scale was set at 96-98 with 100% fluorescent labeled beads. And then the final percentage of EGFP^+^ EVs was normalized against that of the reference beads under the same sensitivity in the fluorescent mode.

#### Automated western blotting of RNP-EV

Total protein extracts from purified RNP-EVs were prepared in the one volume of lysis solution, RIPA buffer (Thermo), followed by 5min sonication, 30s vortex. Protein concentration was then determined by the Micro BCA Protein Assay (Thermo, USA). Protein extracts (100-150ng per lane) were added and separated in the cartridge compatible with Wes Instrument (Bio-Techne, USA). Simple Western was performed and imaged according to the manufacture’s procedure.

#### Transmission electron microscopy

10μL of EVs suspension with 10^9^ particles/mL was vortexed for 30s and then placed on the parafilm. Glow discharged dark side of 400-mesh TEM grid (Electron Microscopy Sciences, USA) was incubated with EV droplet on the parafilm for 15min at room temperature. The TEM grid was then air dried at room temperature. 2% Uranyl acid was used for the negative staining of EVs on the TEM grip for 5min at room temperature. The TEM grid was then washed with deionized water for 10s and air dried for TEM imaging on a Tecnai G2 Sprit TWIN 120kV with UltraScan 1000 (2k x 2k) CCD camera.

#### Immuno-electron microscopy

TEM grid was pretreated with 1% poly-L-Lysine for 15min and then air dried. 10μL of EVs suspension with 10^9^ particles/mL was incubated with primary antibodies (anti-Cas9 or anti-CD81) diluted 1:25 in the PBT buffer containing 1X PBS, 0.1% BSA and 0.01% Tween 20 for 1 hour at 4°C. The resulting solution was then added to TEM grid, dried at room temperature followed by three times washing in 1X PBS and 0.1% BSA. The TEM grid was then incubated in the prepared 1% paraformaldehyde for 5min, followed by two-time washing in PBT buffer for 5min each. The grid was then incubated with secondary antibody (goat anti-mouse and goat anti-rabbit conjugated to 6nm gold) diluted 1:10 in PBT buffer for 1 hour at room temperature, then washed by passing over three droplets of deionized water. The grid was post-stained with 2% uranyl acetate and imaged on a Tecnai G2 Sprit TWIN 120kV with UltraScan 1000 (2k x 2k) CCD camera.

#### RT-qPCR quantification of sgRNA

∼10^8^ EVs or RNP-EVs were firstly treated with 2U Proteinase K (Thermo, USA) for 15min at 37°C and then proteinase K was inhibited by adding 1X Halt protease inhibitor cocktail. 2U RNase A (Thermo) was then added to RNP-EV, native EVs and μDES-EVs, and incubated at 37°C for 15min. The total RNA content from the treated EVs was then isolated by following the manufacturer’s protocol of miRNeasy purification kit (QIAgen, USA). The total EV RNA was then quantified by Ribogreen quantification kit (Thermo, USA). The reverse transcription of gRNA was conducted at 54°C for 30min with AMV Reverse Transcriptase by following the instructions. The resulting cDNA of gRNAs were then directly used for the qPCR quantification with PowerTrack™ SYBR Green Master Mix for QuantStudio (TM)7 Flex System. The final cDNA was aliquoted to four individual qPCR reactions by using the primers in **Table S1**.

### Ototoxicity investigation of LNP and EV

Around 10^8^ EVs and LNPs per HEI-OC1 cell in 96-well plate were used for the biosafety test. Basically, EVs were treated with μDES strategy described before but without the encapsulation of RNP. LNP was also prepared according to the manufacturer’s protocol. The resulting EVs and LNPs were incubated with HEI-OC1 cells in 96 well plate for 72hours. The PrestoBlue reagent (Thermo, USA) was then added to each well to quantify the viability of HEI-OC1, hair-like cell line, based on the manufacturer’s instructions. The fluorescence intensity was subsequently assessed at 560/590 nm using a Cytation 5 microplate reader (BioTek, USA).

### Indel Analysis

#### T7E1 assay

200 ng purified amplicons were denatured and re-annealed by following the manual of GeneArt Cleavage detection kit. The re-annealed amplicons were then incubated with T7 endonuclease I for 30min at 37°C. The final solution was then immediately added to the well of 0.8-2% SYBR safe prestained E-gel and run for 26min in the E-gel running system (Thermo, USA). The gel was then imaged with Typhoon imager using the Cy2 channel. The band intensity was quantified with ImageQuant and Indel % was obtained by using equation: (1-(1-fraction cleaved)^1/2^).

#### Sanger sequencing and ICE analysis

Purified amplicons with more than 200nt flanking sequences were quantified with Cytation 5 and then diluted to the final concentration at 5 ng/μL. The resulting amplicons together with reverse primer (5’-GCGTAGGAGTTGGACTTGATAG - 3’) were then submitted to Sanger sequencing provided by Genewiz. The sequencing data was opened with UGENE software for imaging and visualization of amino acid sequences. The DNA sequencing data were then uploaded to *ICE*, the online quantitative tool of Indel analysis of sequencing trace of CRISPR/Cas9 editing. The Indel% was obtained by comparing the decomposition frequency of CRISPR/Cas9 treated groups to the one without the treatment of CRISPR/Cas9. The same gRNAs were entered into *ICE* for both full length of gRNAs and the corresponding truncated counterparts due to the same cutting site.

#### HTS and CRISPResso2 bioinformatic analysis

The sequences of amplicons listed in **Table S1** were obtained from Next Generation Sequencing. The Indel analysis and mutation types were identified with online genome editing bioinformatic tool, *CRISPResso2* by following the steps published before.^60^ For the analysis of coding sequences in *CRISPResso2*, the exon sequences: 5’-CCTTGGGGAACTTGCTCTCCTCATCGATGAGGGAGATGACGTTCATAGGCGGGTTGGCAATCATG TCCAGTGCTTCCTGGTTGTCAGTGAACTCAATGTGCAGCCAGTCGATGCTCTCCAGGTCGTACTCCC -3’. The sequences of gRNAs and the associated truncated counterparts were used in the *CRISPResso2* analysis.

For allele-specific indel analysis, the wild-type *Myo7a (Myo7a*^*WT*^) and *Myo7a*^*sh1*^ sequences in the EXCEL file were analyzed and sorted out by using LEN and SUBSTITUTE Function by using 5’-CCG-3’ for WT reads. The allele-specific indel % was calculated by using <edited *Myo7a*^*WT*^ sequences/(edited *Myo7a*^*WT*^ sequences and edited *Myo7a*^*sh1*^ sequences).>

For mRNA analysis, CRISPResso2 analysis was performed similarly to the indel analysis mentioned above. To quantify intact, non-edited reads, we used the following sequences: 5’-AGGCCGGAA-3’ and 5’-TTCCGGCCT-3’ for WT reads, and 5’-AGGCGGGAA-3’ and 5’-TTCCCGCCT-3’ for Shaker-1 reads.

### Off target analysis

The off-target prediction by using the off-target algorithm from Integrated DNA technology was performed via entering the Shaker-1 DNA sequence. The potential off targets in **Table S2** based on the gRNA-1 sequence were reported based on the ranking scores. The off targets with the lower score have a higher chance to be edited. Therefore, top 9 off targets were amplified by using the primers listed in the **Table S3**. CRISPResso2 were used for the indel percentage quantification based on the corresponding gRNAs.

### Confocal imaging of cellular uptake

1-2k HEI-OC1 cells were seeded on the cover glass slip which was pre-treated with 1% poly-L-Lysine. 1mL culture media was added to each well and then the cells were incubated to allow the cells to attach overnight. 100μL of 10^10^/mL RNP*^EGFP^-EV, RNP*^EGFP^-LNP were then added to incubate with cells for one hour at 37°C, 5% CO_2_ followed with the removal of the culture media. The treated cells were then washed with 1X PBS without Ca^2+^ and Mg^2+^ (Thermo, USA) twice. The cells were fixed with 4% formaldehyde in 1X PBS for 15min and permeabilized with 0.5% Triton X-100 in 1X PBS for 5min followed with two-time 1X PBS wash. The cells were then incubated with 2% BSA and 22.52 mg/mL glycine in 1X PBS solution for 1 hour to reduce the non-specific binding of primary antibodies. The cells were then washed with 1X PBS twice and incubated with 1:500 anti-EEA1 antibody (Abcam, Cat# ab2900, USA), anti-Cas9 antibody (Genscript, Cat# A01935, USA) 1% BSA mixture overnight at 4°C. The secondary antibodies, Alexa 555 anti-mouse antibody and Alexa 647 anti-Rabbit antibody were then incubated with the cells for 1 hour at 4°C after being washed twice. The cells were stained with nucleus staining dye, DAPI with the final concentration of 0.2μg/mL for 5min. After final 1X PBS washing steps, anti-fade reagent was added to the center of the microscope slides and then the side of the cover slip with treated cells were attached onto the slide. The clear nail polish was added onto the four corners of the glass cover slip to stabilize the glass. The glass slide was then ready for confocal imaging with Zeiss Confocal LSM800 Microscope. The different samples were taken with the same imaging settings including gain power, image resolution et al.

### Animals/Ethics

Shaker 1 mice were purchased from Jackson Laboratory (Bar Harbour, ME, USA) and maintained in a breeding colony. All animal care and procedures were approved by the Institutional Animal Care and Use Committee University of Kansas University Medical Center. Evaluation of EV therapy was tested in Shaker mice aged P4-P180. Outcomes measures included hearing testing using auditory evoked brain stem responses (ABR), and histology/immunofluorescence staining.

### Genotyping

Genomic DNA was extracted from mouse pinna by incubating ear punches with 40μL tissue preparation solution (Sigma, Cat No. T3073) and 40μL extraction solution (Sigma, E7526) at 55°C for 30minutes. This was followed by incubating the mixture at 90°C and for 5 minutes then adding 10μL neutralization solution (Sigma, N3910). Genotype was then determined by PCR assay.The primers 5’-CATGTCCAAGGTCCTCTTCC-3’ forward and 5’-CCTAGAATCAGTGCAGAGCA-3’ reverse were designed to amplify a DNA fragment across the mutation, annealing at 58°C with Dream Taq DNA polymerase (Thermo, EP0702), dNTP mix (Thermo, FERR0192), followed by an MspI digest to give a 328 bp product in homozygous mutants, 189 bp and 139 bp products in wild type, and 328 bp, 189 bp and 139 bp products in heterozygous littermates.

### *In vivo* delivery of RNP-EV to the inner ear

For the surgical procedure, mice were anesthetized with an intraperitoneally administered mixture of ketamine (100 mg/kg), xylazine (5 mg/kg) and acepromazine (2 mg/kg). A dorsal postauricular incision was made and the bulla exposed. For EV delivery, the posterior semicircular canal was exposed as previously described^70^. EV injections consisted of (details on EVs used) in 1μL of volume (delivered using a Hamilton micro syringe with 0.1μL graduations). P4 *Shaker-1* mice were anesthetized with cold as described in reference^71^. The posterior semicircular canal was exposed in the postauricular space and a sharp pick was used to open the canal. EV at a dose of 1μL was injected using a microsyringe as described above. Genotyping was carried out at P21 as described.

### ABR Measurement

ABR thresholds were recorded using the Intelligent Hearing Systems Smart EP program (IHS, Miami, FL, U.S.A.). Animals were anesthetized as described above and kept warm on a heating pad (37°C). Needle electrodes were placed on the vertex (+), behind the left ear (-) and behind the opposite ear (ground). Tone bursts were presented at 4, 8, 16 and 32kHz, with duration of 500μs using a high frequency transducer. Recording was carried out using a total gain equal to 100K and using 100Hz and 15kHz settings for the high and low-pass filters. A minimum of 128 sweeps were presented at 90dB SPL. The SPL was decreased in 10dB steps. Near the threshold level, 5 dB SPL steps using up to 1024 presentations were carried out at each frequency. The threshold was defined as the SPL at which at least one of the waves could be identified in 2 or more repetitions of the recording.

### Immunohistochemistry and histology

The animals were anesthetized with phenobarbital (585 mg/kg) and phenytoin sodium (75 mg/kg) (Beuthanasia®-D Special, Schering-Plough Animal Health Corp., Union, NJ, Canada) i.p. and sacrificed via intracardiac perfusion with 4% paraformaldehyde in 1X PBS. The temporal bones were removed, later the stapes extracted, and the round window was opened. The temporal bones were then postfixed overnight in 4% paraformaldehyde in 1X PBS at 4°C. After rinsing in 1X PBS three times for 30min, the temporal bones were decalcified in 10% EDTA (ethylene diamine tetraacetic acid) for 48h. The temporal bones were rinsed in 1X PBS and embedded in paraffin. Seven μm sections were cut in parallel to the modiolus, mounted on Fisherbrand® Superfrost®/Plus Microscope Slides (Fisher Scientific, Pittsburgh, PA, U.S.A.) followed by the overnight drying process. Samples were deparaffinized and rehydrated in 1X PBS two times for 5min, then washed three times in 0.2% Triton X-100 in 1X PBS for 5min and finally in blocking solution 0.2 % Triton X-100 in 1X PBS with 10% fetal bovine serum for 30min at room temperature. After blocking, specimens were stained with anti-4 hydroxynonenal (R&D Systems MSB 3249) or anti 3-nitrotyrosine, MA1-2770. diluted 1:100 in blocking solution. The tissue was incubated for 48h at 4°C in a humid chamber. After three rinses in 0.2 % Triton X-100 in 1X PBS, immunofluorescent detection was carried out with anti-rabbit IgG (1:50; Alexa Fluor 488 nm; Invitrogen®Inc.). The secondary antibody was incubated for 6h at room temperature in a humid chamber. The slides were rinsed in 0.2 % Triton X-100 in 1X PBS three times for 5min and finally coverslipped with ProLong® Gold antifade reagent (Invitrogen™ Molecular Probes, Eugene, OR, U.S.A.).

### Inner ear tissue dissection for HTS

Both sides of organ of Corti was collected after 6 months since the intratympanic administration of RNP-EVs on the left side while untreated on the right side. DNA and RNA samples were collected by following the manufacturer’s instruction of QIAgen MagAttract HMW DNA Kit and miRNAeasy Mini Kit with DNAse treatment respectively. The resulting nucleic acid samples were then quantified with Agilent Genomic DNA Tape assay and RNA Screen Tape assay respectively. cDNA library of mRNA and full transcripts were prepared by using the amplicons for mRNA of *Myo7a* gene shown in the **Table S1**.

### Statistical Analysis

Statistical analyses were performed using GraphPad Prism 9.0.0. Data were presented as biological replicates and were shown as means±SEM or range as indicated in the corresponding figure legends. Meanwhile, sample sizes and the statistical tests used were described in detail in the corresponding figure legends. Unpaired Student’s t test and one/two-way analysis of variance (ANOVA) tests were performed following the appropriate post hoc test when more than two groups were compared. For all analyses, P<0.05 was assigned as statistically significant.

### *In silico* analysis of principal indel variants

Based on the indel frequency from *in vitro* experiments and CRISPResso2 analysis, we first sorted and grouped the new alleles into the 5 principal mutation types. To understand the potential impact of the novel variants from Cas9-mutagenesis on the biological systems, our strategy is obtaining and understanding the associated changes at the protein level. To this end, the Cas9-mutagenesis-derived mRNA sequences that can be transcribed from the indel alleles were entered to Expasy translation tool (https://web.expasy.org/translate/) and the potential polypeptide sequences were collected. The domain selection is based on the features of the unconventional myosin-VIIa (P97479) UniProt database (https://www.uniprot.org/uniprotkb/P97479/entry). Based on the loci where the mutation identified in the annotated domain region, myosin motor domain, we selected residue 401-561 for the further characterization and analysis. The name entry is based on their deletion. The in-frame mutants were characterized with Alphafold2 to predict the structures and obtain PDB files (https://colab.research.google.com/github/sokrypton/ColabFold/blob/main/AlphaFold2ipynb, accessed on X, October 2023) with the following parameters: num_relax=1; template_mode=none; rank_num=3 while the other as default. The resulting PDB files were used for the binary structure alignment and color by predicted pLDDT value by using command line <spectrum b, yellow_orange_cyan_deepblue, minimum 57, maximum 98> in Pymol software. To further understand that the possible polypeptides sequences translated from the all possible reading frames within the new alleles, we then used NCBI BLASTX to explore the potential novel proteins and the associated functions from the top10 DNA sequences containing indels from 1450nt to 1600nt by aligning to the NCBI protein reference database. The final results were downloaded and shown in Excel file.

### Reporting Summary

Further information on research design is available in the Nature Research Reporting Summary linked to this article.

## Data availability

The main data supporting the results of this study are available within the paper and its Supplementary Information files. Source data for the figures in this study are available from the corresponding author upon reasonable request. NGS sequencing data are available at NCBI BioProject ID: PRJNA1041447

## List of Supplementary Materials

Fig S1 to S9 for supplementary figures

Tables S1 to S3 for supplementary tables

Supplementary note I-II

References *1-8*

## Acknowledgments

The authors would like to thank Interdisciplinary Center for Biotechnology Research (ICBR) core facility at the University of Florida, and imaging core facility support from University of Kansas Medical Center.

## Funding

We acknowledge funding support to Dr Mei He as follows:

National Institutes of Health grant 1R35GM133794 (MH)

Cystic Fibrosis Foundation grant HE21I0 (MH)

University of Florida Health Cancer Center GU pilot (MH)

## Author contributions

Conceptualization: Mei He, Hinrich Staecker

Methodology: Mei He, Hinrich Staecker, Xiaoshu Pan, Peixin Huang, Athanasia Warnecke

Investigation: Xiaoshu Pan, Peixin Huang, Samantha Ali, Tarun Hutchinson, Nina Erwin, Zachary Greenberg, Zuo Ding, Yanjun Li, Natalia Fernadez, Bryan Renslo

Visualization: Mei He, Hinrich Staecker, Xiaoshu Pan, Peixin Huang

Funding acquisition: Mei He, Hinrich Staecker

Project administration: Mei He, Hinrich Staecker

Supervision: Mei He, Hinrich Staecker

Writing – original draft: Mei He, Hinrich Staecker, Xiaoshu Pan

Writing – review & editing: Mei He, Hinrich Staecker, Xiaoshu Pan

## Competing interests

Authors declare that they have no competing interests.

## Data and materials availability

All data are available in the main text or the supplementary materials.

## Notes

### Competing Interest Statement

The authors have declared no competing interest.

### Summary of Updates

More animal work was done to solidify the conclusions and updated in this version in Figure 4 and Figure 5.

