## Supplementary Materials for "CRISPR-Cas9 Engineered Extracellular Vesicles for the Treatment of Dominant Progressive Hearing Loss"

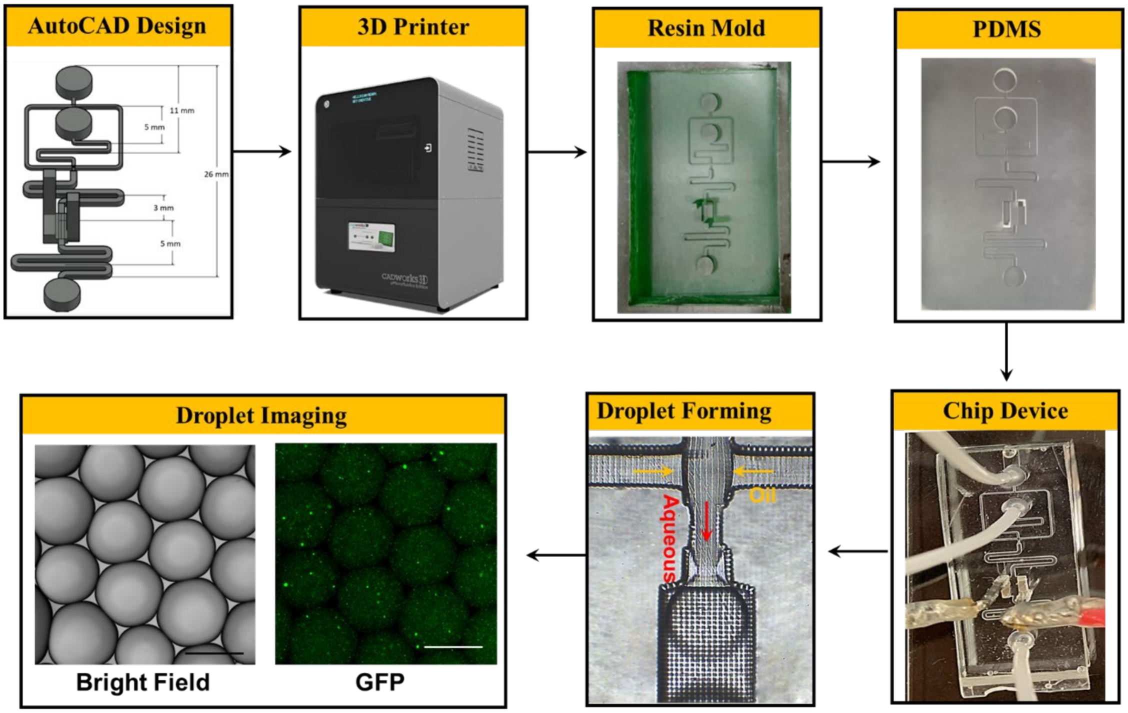


**Supplementary Figure 1.** Workflow for fabricating μDES microfluidic device via simple and low-cost 3D printing and PDMS molding. The characterization of resulting droplets using microscopic imaging showed uniformly sized droplets in ~1000 μm. The droplets encapsulated with fluorescein conjugated Polystyrene latex reference standard (Applied Microspheres, Netherlands) are for mimicking EVs encapsulation in droplets for electroporation. Scale bar: 1000μm.

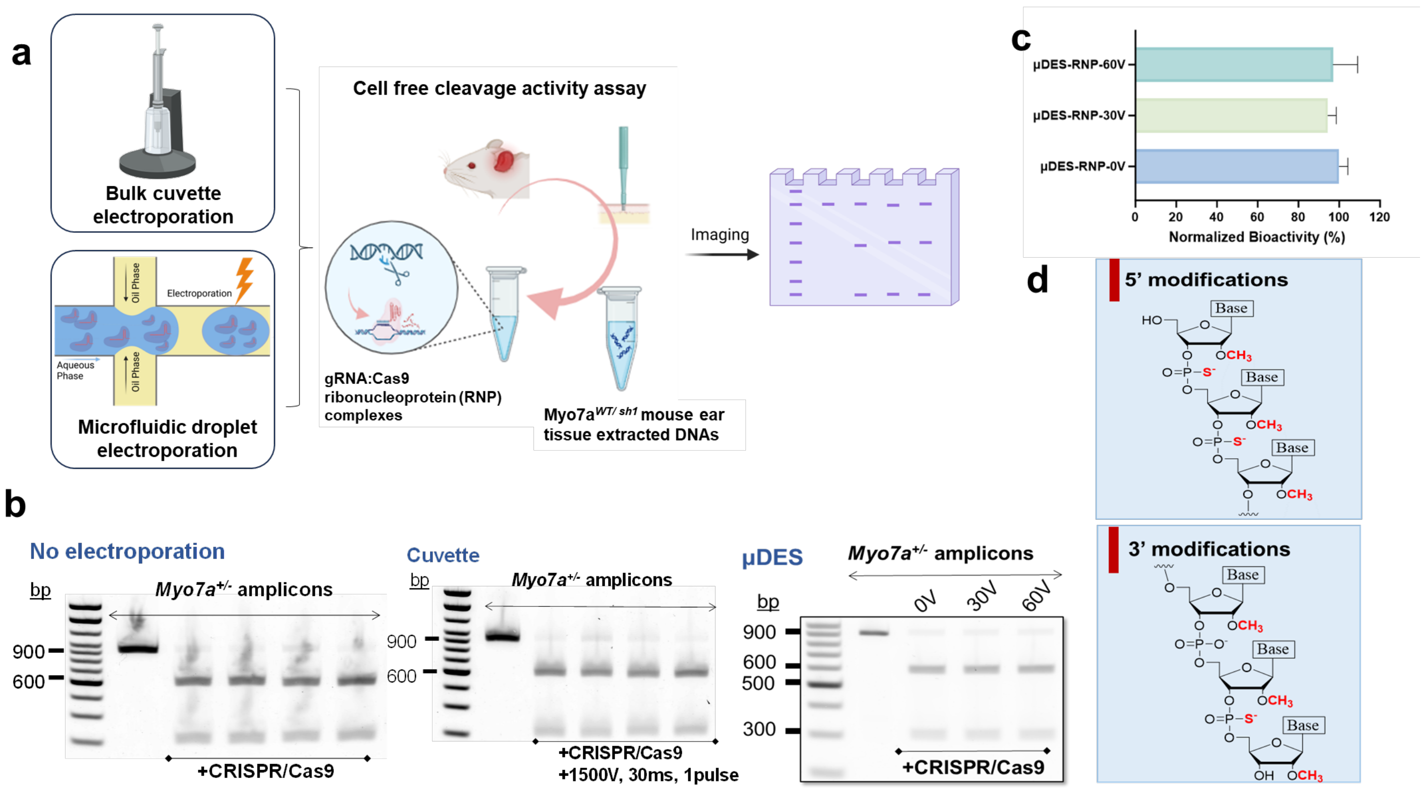


**Supplementary Figure 2. a,** Schematic illustration of Cell free cleavage activity assay to evaluate the activity of gRNA:Cas9 ribonucleoprotein (RNP) complexes under different electroporation conditions. The PCR amplicons were amplified by using genomic DNA isolated from the Myo7a $^{WT/sh1}$ mouse ear punch as the template. **b,** DNA gel electrophoresis to evaluate the cleavage fragments from cell-free cleavage assay under conditions (left) without electroporation on RNPs, (middle) bulk cuvette electroporation, (right) microfluidic droplet electroporation. **c,** Quantification of bioactivity on CRISPR/Cas9 RNPs showed little or no influence on the overall function after µDES treatment, which is comparable to the native status of RNPs while editing Myo7a $^{WT/sh1}$. **d,** The schematic illustration of chemical modifications on the guide RNA sequences used in the study.

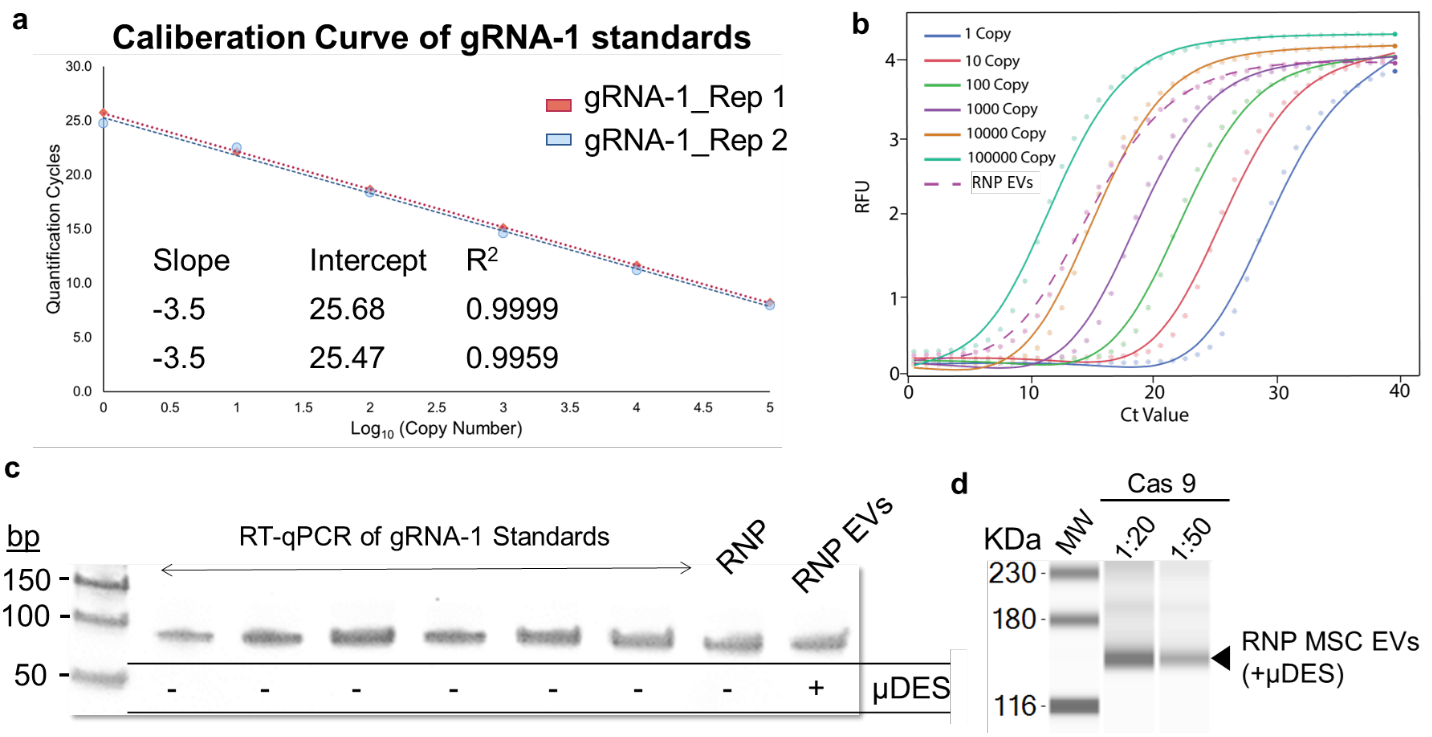


**Supplementary Figure 3. a,** RT-qPCR quantification of gRNAs using calibration Standard curve to quantify the gRNAs in the total RNA content purified from RNP EV lysates. Each data point represents 2-3 technical replicates, and each curve data set is from the independent experimental replicates. **b,** Amplification plot of RT-qPCR quantification of gRNA standards with known copy numbers. The µDES produced RNP EV group showed their quantitation is within the range of the serial dilution of gRNA standards. **c,** Electrophoretic assay of the gRNA-1 amplicons after RT-qPCR experiments showed the amplicons' size which is the same as that from RNPs or µDES produced RNP EV group, indicating the specific identification of gRNAs under different experimental conditions. **d,** The Western blotting analysis showed RNP loading into MSC EVs. Lane 1: MW protein markers; Lane 2: 1:20 anti-Cas9 antibody against lysate from 0.025mg/mL µDES produced RNP EVs; Lane 3: 1:50 anti-Cas9 antibody against lysate from 0.025mg/mL µDES produced RNP EVs.

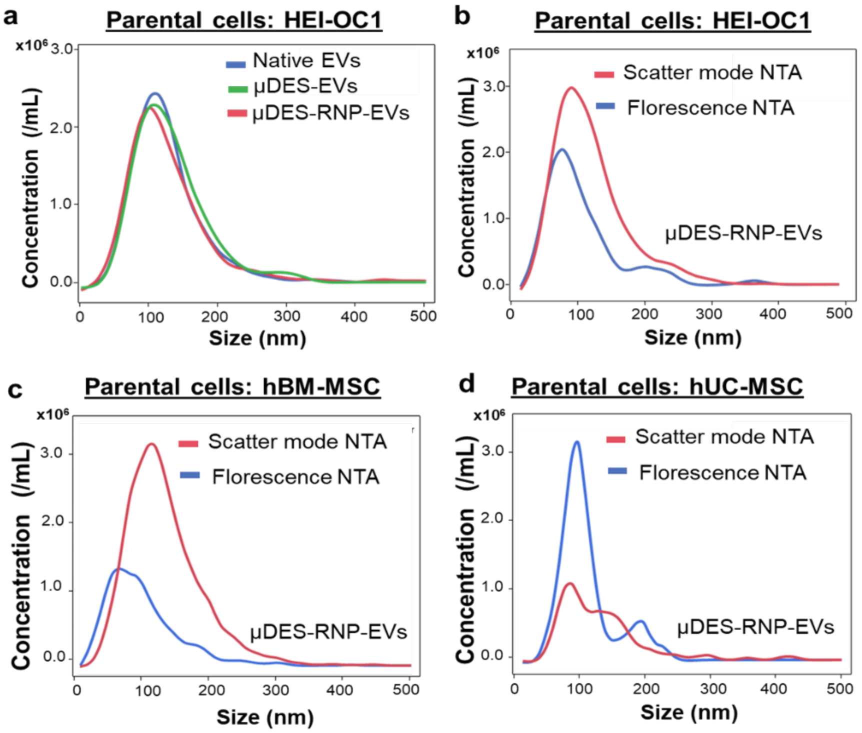


**Supplementary Figure 4**. Nanoparticle tracking characterization of the size of RNP-loaded EVs. **a,** EVs secreted from HEI-OC1 cells (native EVs) were used for microfluidic droplet electroporation (µDES) with and without the fluorescent cargo RNP$^{*EGFP}$. The NTA measurements were performed under scattering mode. **b,** HEI-OC1 cells secreted EVs were employed to load the cargo RNP$^{*EGFP}$ to them via µDES. The NTA measurements were conducted using both scattering mode and fluorescence mode (Ex488nm/Em505nm). **c,** Human bone marrow-derived MSC cell (hBM-MSC) secreted EVs were used for to load cargo RNP$^{*EGFP}$ to them. The NTA measurements were performed with both scattering mode and fluorescence mode (Ex488nm/Em505nm). **d,** Human umbilical cord-derived MSC (hUC-MSC) cell secreted EVs were used for µDES to load cargo RNP$^{*EGFP}$. The NTA measurements were performed with both scattering mode and fluorescence mode (Ex488nm/Em505nm).

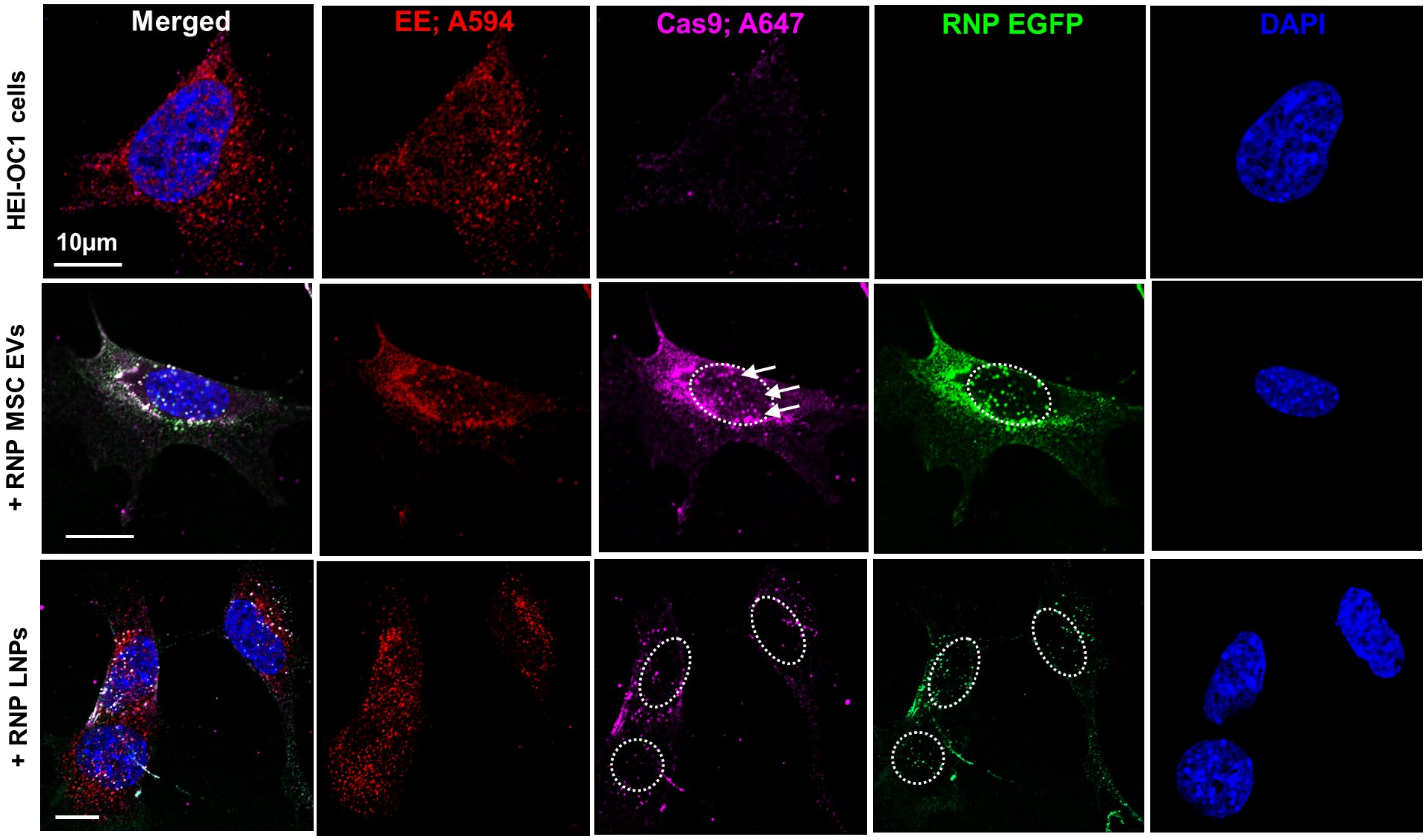


**Supplementary Figure 5.** Confocal imaging analysis of HEI-OC1 hair cells for uptaking particles in one-hour following dosing with RNP*$^{EGFP}$ LNPs (LNP-102, Cayman Chemical) and RNP*$^{EGFP}$ MSC-EVs in ~$10^9$ particles. The white arrow indicates the cytoplasmic release and gradual entry into the nucleus. Scale bar = 10 µm. Confocal imaging on cellular uptake of RNP EVs and RNP LNPs respectively within one hour. The bone marrow derived MSC EVs with loaded RNPs and RNP LNPs were incubated with HEI-OC1 cells individually ($10^6$ Particles/cell) for one hour.

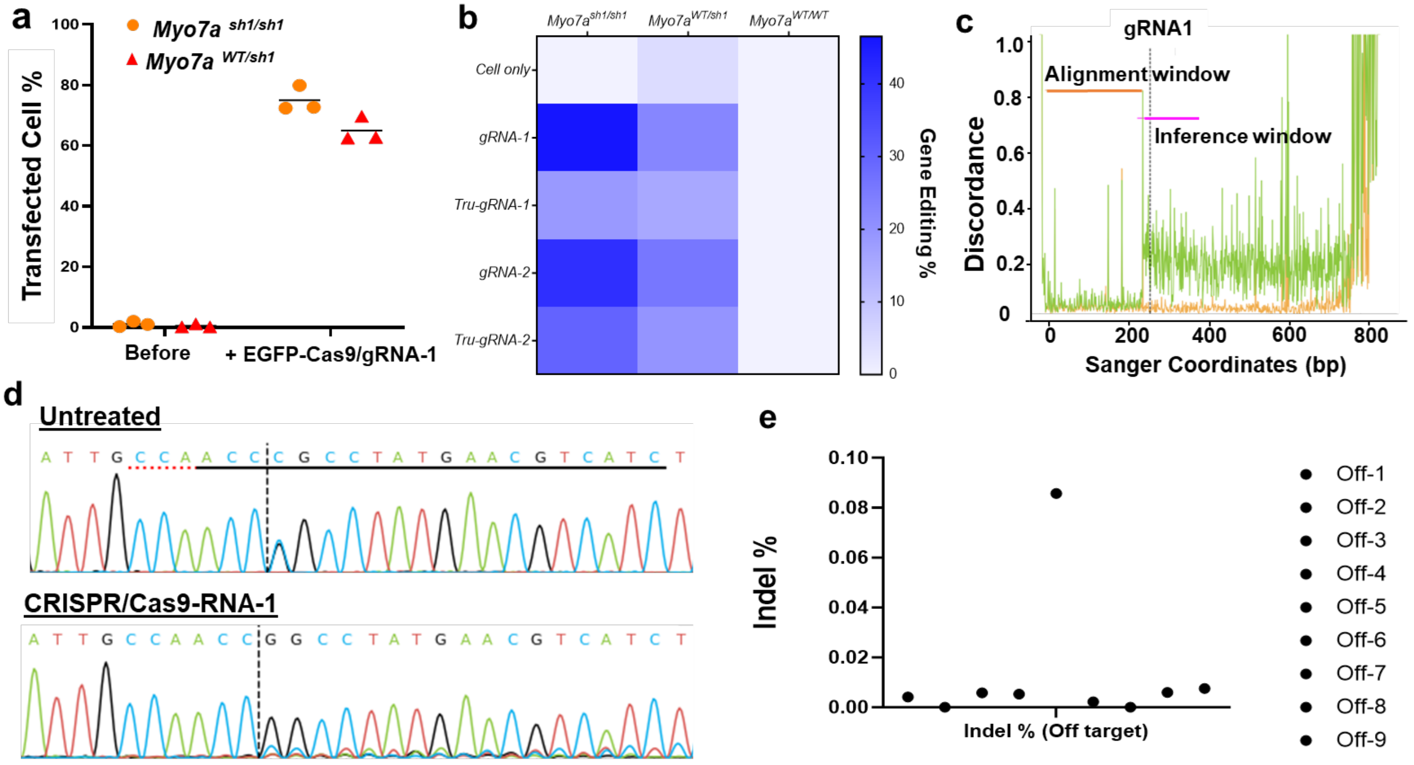


**Supplementary Figure 6**. Characterization of allele-specific editing *in vitro* using Shaker-1 mouse ear tissue derived fibroblast cells. **a,** Flow cytometric quantification of transfected cells using traditional bulk cuvette electroporation of EGFP tagged Cas9 which was complexed with gRNA-1 to form RNPs. The fibroblast cells are from both heterozygous Shaker-1 (*Myo7a*$^{WT/sh1}$) mice and homozygous Shaker-1 mice (*Myo7a*$^{sh1/sh1}$). **b,** Heat map of gene editing % from using different gRNAs: gRNA-1, Tru-gRNA-1, gRNA-2, Tru-gRNA-2. By using genomic mutation detection assay, T7E1, the calculation of gene editing % is based on the equation: gene modification = 100 x (1 – (1- fraction cleaved)1/2). The *in vitro* indel assays indicated the good editing ability of CRISPR systems against *Myo7a sh1* mutants while having little editing effect on *Myo7a WT*. **d,** Sanger sequencing chromatograms of *Myo7a* amplicons from the fibroblast cells with and without treatment of CRISPR/Cas9-gRNA-1. **e,** Off target analysis of Top 9 DNA sequences listed in the Table S2. The potential indel percentage is lower than 0.01% at the DNA level in vitro.

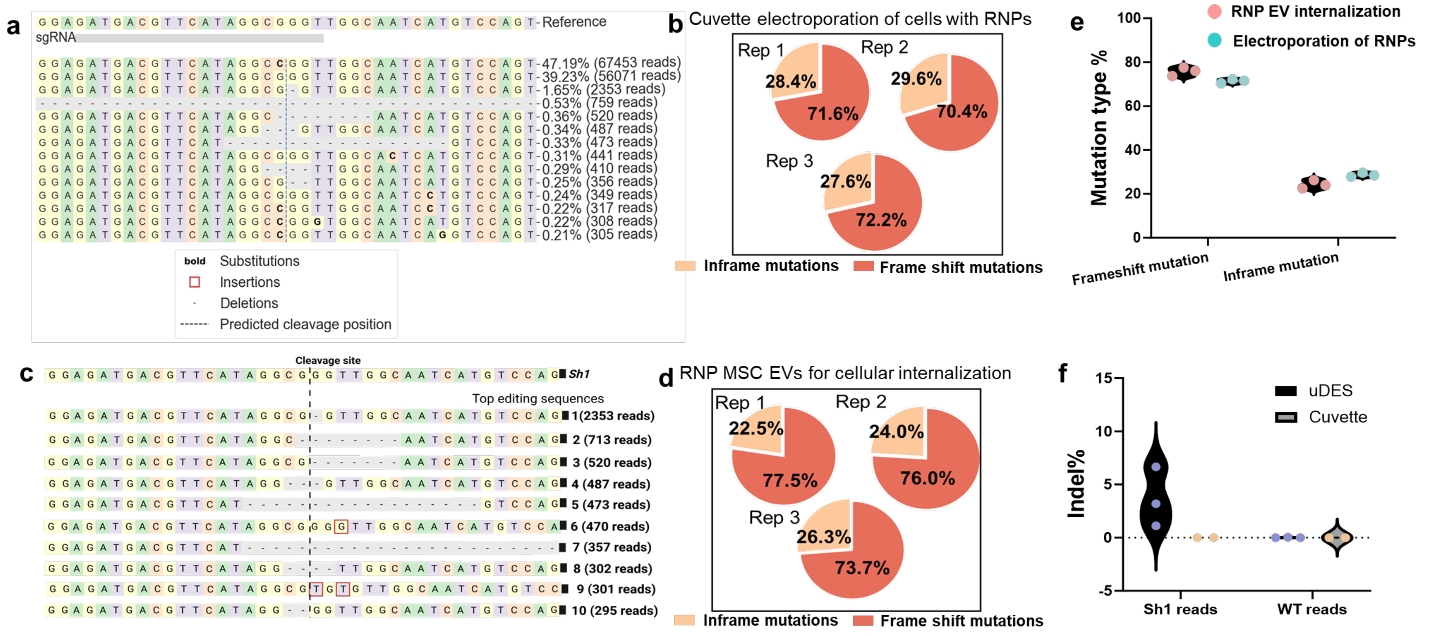


**Supplementary Figure 7**. **a,** Indel frequency table from CRISPResso2 showed that µDES produced RNP MSC EVs can effectively disrupt *Myo7a*$^{sh1}$ allele in the *Myo7a*$^{WT/Sh1}$ fibroblasts. **b,** Comparison of editing continuity from the different biological replicates in the pie chart regarding the mutation types demonstrated by direct electroporation of RNP to cells which had similar in-frame and frame shift mutation percentages. **c,** Summary table of the notable indel alleles from three different biological replicates of fibroblast cells treated with RNP MSC EVs which showed that the most abundant indel allele is the single base deletion. The number of reads is from individual biological replicate. **d,** Comparison of editing continuity regarding the mutation types from three different biological replicates demonstrated by cellular internalization of RNP MSC EVs, which had similar in-frame and frame shift mutation percentages. **e,** Comparison of two mutation types from direct RNP electroporation and EV-mediated RNP delivery which displayed high similarity of desired mutation types, suggesting stable and consistent genome editing behavior mediated by EV delivery. **f,** Quantification of indel percentage demonstrated that RNP MSC EVS led to higher indel percentage from µDES system than that from cuvette bulk Neon electroporation system. The dosing concentration is $10^7$ EVs per fibroblast cell.

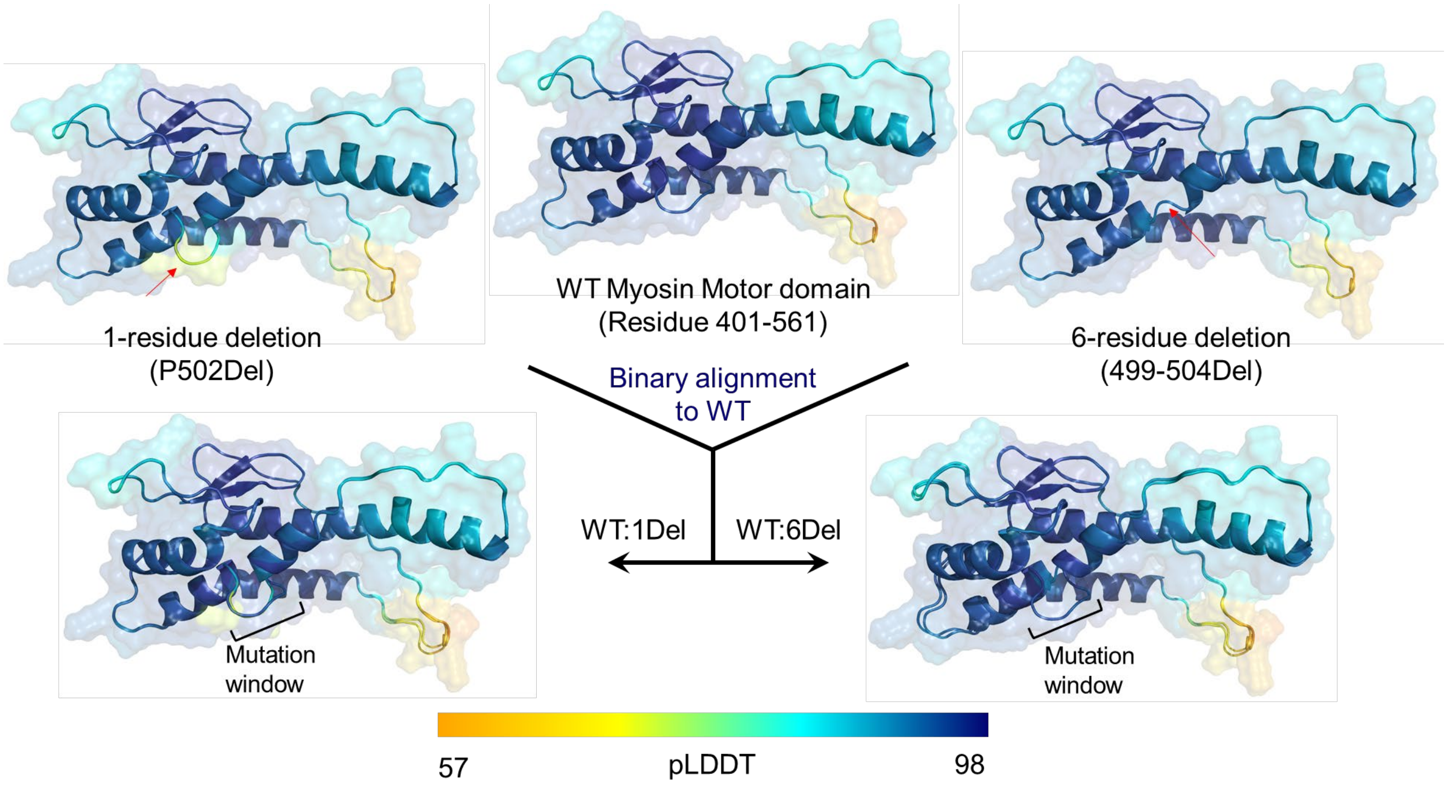


**Supplementary Figure 8.** Per-residue confidence scores (pLDDT) from AlphaFold2 for 1-residue deletion and 6-residue deletion were mapped onto the highest scoring structures from the AlphaFold2 protocol.

**Supplementary note I of Supplementary Figure 8:** Canonical *Myo7a* protein is composed of five major domains, myosin motor, MyTH4, FERM, IQ, and SH3 as reported before.[1] The natural shaker-1 variant replaces Arg with Pro at the 502nd residue within the myosin motor domain of *Myo7a*.[2] The *in vitro* NGS results (Fig. 3f) showed that two notable sequences represented in-frame mutation which was induced by Cas9-mutagenesis. Considering that no reported evidence to indicate the exon splicing enhancer or any functional splicing events within this cleavage site,[3] the protein assembly containing in-frame mutation may not be interfered thereby generating the full length of the *Myo7a*-like protein variants. To understand the potential impact of in-frame mutations on protein structures, we employed Alphafold2 pipeline with relaxation to characterize the structural similarity to the residue 401-561 in the wild-type motor domain where harbored shaker-1 mutation along with Cas-induced mutants.[4] The resulting three structures showed overall very high per-residue confidence scores, annotated by pLDDT, especially in the structured helical regions which were well-documented in multiple reports.[5] The 1-residue deletion domain exhibited relatively lower pLDDT scores in the unstructured loop region (red arrow in Supplementary Figure 8) when compared to the high mean pLDDT value in the structured regions which are still around 80, indicating high confidence. Jointly considering the loop region was mostly tolerant to the indel in general[5-6] and the basis of AlphaFold2's prediction[4-5, 7], we expect that 1-residue deletion on the loop region is unlikely to reduce significant structural integrity compared to WT domain shown in the Supplementary Figure 8. Similarly, the pLDDT of 6-residue deletion of the loop region also showed high tolerability to the deletions and high foldability, while the binary alignment to WT exhibited slightly different positioning 6-residue deletion domain in Supplementary Figure 8. The emerging evidence showed that the copy numbers of WT *Myo7a* can play an important role in the Shaker-1 mouse models.[1, 8] By reducing percentage of Shaker-1 allele by Cas9-mutagenesis, WT *Myo7a* allele to Shaker-1 allele is notably enhanced *in vitro* and *in vivo*. Therefore, the functional WT *Myo7a* proteins can dominate over the possible variants from potentially translatable alleles. Such domination is greatly beneficial for the prevention of hearing impairment. Though it is still challenging to predict the gradual or drastic shifts on the function of final in-frame mutants based on the computational toolkits,[5] the residual expression level and the associated high structural integrity render potentially limited impacts of in-frame mutants on the biological function such as hearing ability.

Moreover, we then explore the amino acid sequences derived from the top 10 allele sequences, ranked by their abundance by using NCBI Protein Reference Sequences and NCBI BLASTX algorithm. Upon analysis, we did not observe any significant alignments that would generate harmful protein hits within the current amplicon size of our sequencing design. Consequently, it is envisaged that the RNP delivery system may not compromise the integrity of the hearing system, from the DNA level to the protein level, as indicated by our bioinformatic and state-of-art protein modeling analysis.

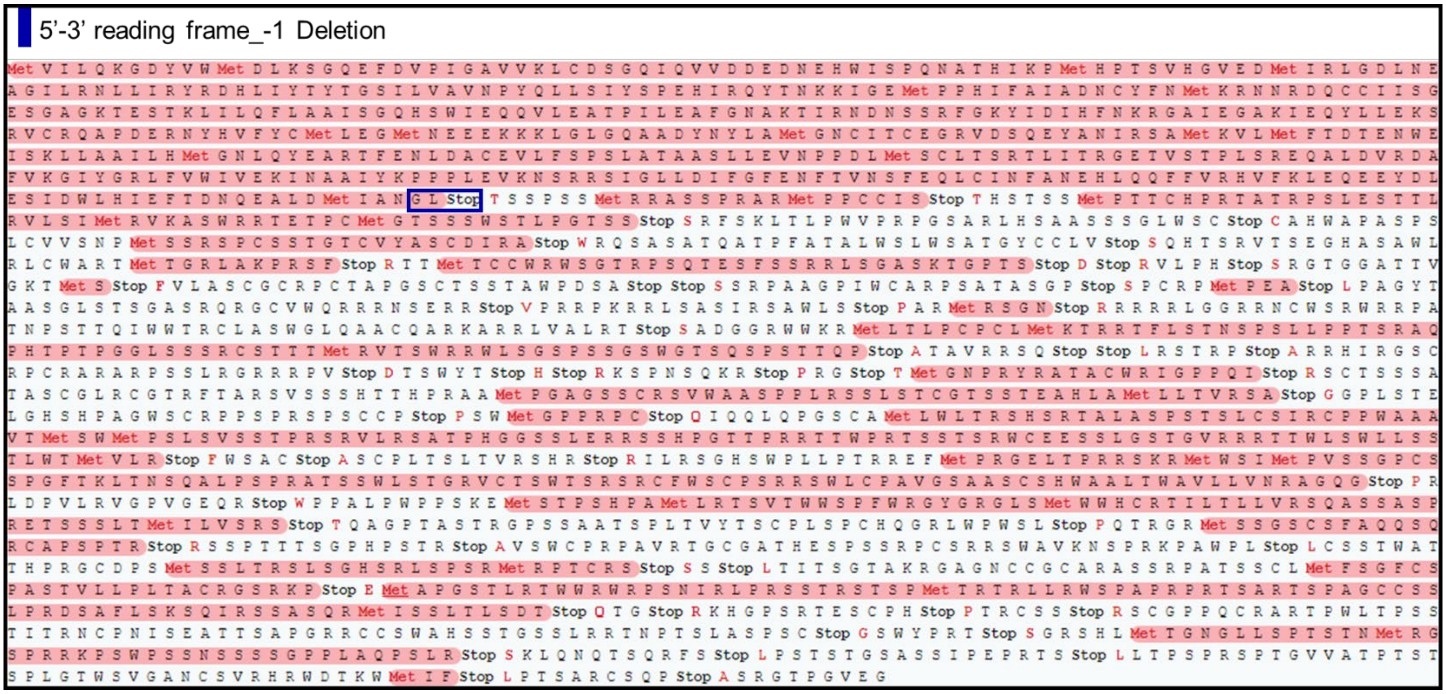


**Supplementary Figure 9.** The analysis of frameshift mutations at the protein level revealed the emergence of premature stop codons, implying potential disruption in the complete protein assembly of the *Myo7a* variant resulting from Cas9-mediated mutagenesis. Specifically, the polypeptide sequence in red showed that the translation from around 70-80% of indel alleles was prematurely terminated at the 504$^{rd}$ residue following the subtitution of residues at position 502-503. (Open reading frames are highlighted in red. The window of mutagenesis window was highlighted in blue square.)

**Supplementary note II of Supplementary Figure 9:** CRISPResso analysis of sequencing results revealed that the most abundant sequences (>1%) with indels in the entire cell pools were located in the 13$^{th}$ exon site after the treatment of gRNA-1:Cas9 RNP. To date, to our best knowledge, there is no rare splicing site reported within the 13$^{th}$ exon region.[3] Therefore, it is reasonable to infer that the most likely protein sequences can directly result from the indel allele containing indels in the coding region, without taking into account RNA splicing or exon skipping. We then altered the mRNA sequence of *Myo7a* gene based on the editing outcomes. Subsequently, the sequences were inputted to web-based Expasy translation tool to predict the potential protein sequences. In the case of single-base deletion frameshift mutation, the polypeptide sequences were fragmented due to the introduction of premature stop codons, suggesting the disruption of translation for functional *Myo7a* variants. The resulting fragment, comprising approximately 503 amino acids, is notably shorter than the full-length *Myo7a*$^{Sh1}$ or *Myo7a*$^{WT}$, which consist of 2215 amino acids. Therefore, it is less likely to that this truncated variant retains functional attributes, rendering it less possible for introducing new *Myo7a*-like variants associated with deafness.

**Table S1.** Sequences of gRNAs, PCR primers used for analysis of on-target editing efficiency with the T7E1 assay, next generation sequencing and quantitative PCR. * Indicates the chemical modifications on the first three bases of gRNA sequences.

| Name | Sequences (5'- 3') |
|---|---|
| gRNA-1 | G*A*U* GAC GUU CAU AGG C**G**G GU |
| Tru-gRNA-1 | G*A*C* GUU CAU AGG C**G**G GU |
| gRNA-2 | A*G*G* GAG AUG ACG UUC AUA GG |
| Tru-gRNA-2 | G*A*G* AUG ACG UUC AUA GG |
| *Myo7a*_FP_T7E1 | GAG GGA ACA GAG TGG CTA TTA C |
| *Myo7a*_RP_T7E1 | GCG TAG GAG TTG GAC TTG ATA G |
| *Myo7a*_FP_NGS | ACA CTC TTT CCC TAC ACG ACG CTC TTC CGA TCT CCC AGG TCA AGC CAA TTC TAT |
| *Myo7a*_RP_NGS | GAC TGG AGT TCA GAC GTG TGC TCT TCC GAT CTC TTC GAG CAG CTC TGC ATT A |
| mRNA_*Myo7a*_FP | CAT CGA CTG GTT GCA CAT TG |
| mRNA_*Myo7a*_RP | TGA GCT TGT GCT GTG AGT T |
| 18s rRNA_Mouse_FP | GCA ATT ATT CCC CAT GAA CG |
| 18s rRNA_Mouse_RP | GGC CTC ACT AAA CCA TCC AA |
| *MYO7A*_Mouse_FP# | CCG CCT ATG AAC GTC ATC TC (-1 bp) |
| *MYO7A*_Mouse_RP# | TGA GCT TGT GCT GTG AGT T (+79 bp) |

#Shaker-1 mutation is annotated as "0". The direction to 5' end of mRNA of MYO7A is annotated as "-" while 3' end as "+". For instance, 5'- AAC CGG CCT A -3'. Fwd primer was counted from 5' end while Rev primer from 3' end.
-2 +2

**Table S2.** All primers are designed for the NGS-based off-target analysis *in vitro*.

| Name | Sequences (5'- 3') |
|---|---|
| Off-1_FP | CCC TTC CAC CCT TTC CTA ATC |
| Off-1_RP | AGG CTC CTG TGT ACT TCT CT |
| Off-2_FP | TCA AAG AGT CAA ACC CGA ACT |
| Off-2_RP | TTC AGC AAG GCA GCA AGA |
| Off-3_FP | CTG GTG GCA ATG TGC AAA TAA |
| Off-3_RP | CCA TGA GCA CTG TTC ACT ATC T |
| Off-4_FP | GCC GAG AAT AAC CTG GAA AGA |
| Off-4_RP | CCA TCA TTG AGG AAA GCC AAA G |
| Off-5_FP | CAC TAG GTG TGT GAG TGA GTT C |
| Off-5_RP | AGC TGC TAT TGC TGG TTA AGT |
| Off-6_FP | TTG GTG ACC ACC AGT GTA TG |
| Off-6_RP | CTC TAC TGT CCC TGA GCT AGA A |
| Off-7_FP | GGC AGC TTT CCA TCA GAA ATG |
| Off-7_RP | TGG CAT CCT GGA ACT CTA ATG |
| Off-8_FP | TGG CAT CCT GGA ACT CTA ATG |
| Off-8_RP | CAG CTT ACG GAT GGC AGA TAC |
| Off-9_FP | CCC TCA TAA GCA CCA CAT ACA |
| Off-9_RP | GGA ACC TTA CCT GCC TTA GAA T |

**Table S3.** Computationally reported Top 18 off targets and the associated scores, gene, locus and the associated gRNA sequences

| Name | Aligned gRNA Sequences | Off target score (IDT) | Number of Mismatches | Gene | Locus |
|---|---|---|---|---|---|
| Off-01 | GTTAACG-TCATAGGCAGGT | 10 | 4 | | chr18:+78063509 |
| Off-02 | CATGTACTTTCATAGGCAGGT | 16 | 4 | | chr11:-32840609 |
| Off-03 | AATG-CTTTCACAGGCGGGT | 19 | 4 | | chr3:+129557655 |
| Off-04 | AATGCCATTCATAGGCAGGT | 23 | 4 | | chrX:+153653923 |
| Off-05 | CATGACATTCATAGGTGGGT | 27 | 3 | | chr18:+30939732 |
| Off-06 | ATTGACATTC-TAGGCGGGT | 28 | 4 | | chr17:+60892023 |
| Off-07 | GATGCTCGTTCATAGGCAGTT | 32 | 4 | | chr6:+95391158 |
| Off-08 | GGTGACATTCATAAGCAGGT | 33 | 4 | | chr15:+89398713 |
| Off-09 | GATGACATGTCATAGGAGGGT | 34 | 3 | | chrX:-36743194 |
| Off-10 | GTTTACTTTCATAGGTGGGT | 34 | 4 | | chr6:+47129576 |
| Off-11 | GAGGAAATTCATAGACGGGT | 36 | 4 | | chr10:+16485319 |
| Off-12 | GATGACGTTCATAGGCCGGT | 39 | 1 | MYO7A | chr7:+98092439 |
| Off-13 | GGTGTCGTTCCTAGGCAGGT | 41 | 4 | | chr5:-45374917 |
| Off-14 | GATAAC-TTCATAGGTGGGT | 43 | 3 | | chrX:+11299227 |
| Off-15 | GATAAC-TTCATAGGTGGGT | 43 | 3 | | chrX:-9572422 |
| Off-16 | GATAAC-TTCATAGGTGGGT | 43 | 3 | | chrX:+11318231 |
| Off-17 | GATAAC-TTCATAGGTGGGT | 43 | 3 | | chrY:-2720748 |
| Off-18 | GATAAC-TTCATAGGTGGGT | 43 | 3 | | chr2:-17996948 |
